# Redox state and cellular uptake of copper is regulated by N-terminus of human Copper Transporter-1

**DOI:** 10.1101/2021.08.03.454867

**Authors:** Sumanta Kar, Samarpita Sen, Saptarshi Maji, Ruturaj, Rupam Paul, Sohini Dutt, Basudeb Mondal, Enrique Rodriguez-Boulan, Ryan Schreiner, Durba Sengupta, Arnab Gupta

**Affiliations:** Department of Biological Sciences, Indian Institute of Science Education and Research Kolkata, Mohanpur, West Bengal 741246, India; Department of Chemical Sciences, Indian Institute of Science Education and Research Kolkata, Mohanpur, West Bengal 741246, India; Department of Ophthalmology, Margaret Dyson Vision Research Institute, Weill Cornell Medicine, New York, NY10021, United States; CSIR-National Chemical Laboratory, Dr. Homi Bhabha Road, Pune, Maharashtra 411008, India

**Keywords:** CTR1, SLC31A1, Copper homeostasis, polarized epithelia, MDCK

## Abstract

Copper(I) is essential for all life forms. Though Cu(II) is most abundant state in environment, its reduction to Cu(I) is prerequisite for bio-utilization, by a mechanism that is uncharacterized. We show that in human Copper Transporter-1, two amino-terminal methionine-histidine clusters and neighbouring aspartates distinctly binds Cu(II) and Cu(I) preceding its import. The endocytosis of hCTR1 from basolateral membrane of polarized epithelia to Common-Recycling-Endosomes is dependent on copper reduction and Cu(I) coordination by methionines. The transient binding of both Cu(II) and Cu(I) during the reduction process facilitated by aspartates acts as another crucial determinant of hCTR1 endocytosis. Mutating ^7^Met-Gly-Met^9^ and Asp^13^ abrogates copper uptake and endocytosis that is correctable by reduced and non-reoxidizable Cu(I). Histidines clusters are crucial for hCTR1 functioning at limiting copper. Finally, we show that two N-terminal His-Met-Asp clusters exhibit functional complementarity in regulating Cu(I)-induced hCTR1 endocytosis. We propose a mechanistic model where His-Met-Asp residues of amino-terminal hCTR1 coordinates copper and maintains its reduced state crucial for uptake.

## Introduction

Copper is a micronutrient essential for all eukaryotic organisms as it plays a crucial role in coordinating different physiological activities of cells(Fritsch et al., 1988; Linder, 1991; Labbé et al., 1999; Puig et al., 2002). Copper shuttles between its two primary oxidation states, Cu(I) and Cu(II). Though it participates in physiological processes, the cuprous ion Cu(I) is unstable in the oxidizing environment; Cu(II) is the most abundant oxidation state in hydrophilic and oxidizing environments(Bertinato, 2015). In yeasts, copper is first reduced from Cu(II) to Cu(I) by cell surface reductases Fre1/Fre2 prior to uptake(Hassett and Kosman, 1995; Georgatsou et al., 1997; Yun et al., 2001). However, the mechanism of Cu(II) reduction in mammalian cells is not clearly understood. In this study, we, for the first time, show that the human Copper Transporter-1 (hCTR1), besides importing copper, also plays a crucial role in maintaining the Cu(I) redox state that renders the metal bioavailable for physiological utilization in cells.

CTR1 (SLC31A1) is the only high-affinity plasma membrane copper importer that has been known to date in mammalian cells(Zhou and Gitschier, 1997; Lee et al., 2002b). CTR1 is the primary member of the CTR family consisting of six known members (CTR1-6), with at least one member found in all eukaryotic life forms(Mandal et al., 2020). The human CTR1 (hCTR1) is a small protein of 21 kDa, consisting of 190 amino acids. It exists as a homo-trimeric integral membrane protein with each monomer consisting of an extracellular amino-terminal (N-term), three transmembrane (TM) domains, and a small intracellular cytosolic tail(Gupta and Lutsenko, 2009; Eisses and Kaplan, 2002; De Feo et al., 2009). The extracellular N-term shows high sequence variability among species. hCTR1 N-term is 67 amino acids long and is predicted to be unstructured(De Feo et al., 2009). The amino terminus of hCTR1 also harbours N-linked (Asn^15^) and O-linked glycosylation (Thr^27^). O-linked glycosylation at Thr-27 is necessary to prevent proteolytic cleavage that removes approximately half of the N-term of hCTR1(Maryon et al., 2007a).

In unpolarized epithelial cells, e.g., HEK293T, hCTR1 localizes on the plasma membrane in basal or copper limiting conditions(Petris et al., 2003). In high copper treatment, as a self-regulatory mechanism to limit copper import, hCTR1 endocytoses in vesicles and accumulates in early sorting and recycling endosomes marked by Rab5 and EEA1(Clifford et al., 2016). Using live-cell imaging, Clifford et al. demonstrated that, upon removal of extracellular copper, hCTR1 recycles back through the Rab11 dependent pathway(Clifford et al., 2016). Using another unpolarized model, i.e., HeLa cells, Curnock and Cullen have further shown that the retromer complex regulates plasma membrane recycling of the protein and prevents it from entering the lysosomal degradation pathway(Curnock and Cullen, 2020). However, localization of hCTR1 in polarized epithelia has been a field of debate with several contradictory reports. Data from Thiele group has shown in mouse intestine that hCTR1 localizes at the apical membrane and is responsible for luminal copper uptake(Nose et al., 2006, 2010). On the other hand, Zimnicka *et al*. demonstrated that hCTR1 localizes on the basolateral surface of kidney epithelial cell model, MDCK, enterocyte models Caco-2, as well as a model for intestinal crypt cells, T84(Zimnicka et al., 2007). This observation was also reproduced in mouse intestinal sections(Zimnicka et al., 2007). A clearer understanding of hCTR1 localization in the polarized epithelial cell is warranted as its crucial extracellular N-terminal domain will be exposed to two completely different extracellular environments found in luminal (apical) vs. blood (basolateral) side that will possibly influence copper availability, mechanism of copper-binding and uptake by the protein.

*In-vitro* as well as *in-vivo* studies have shown that the N-terminal domain can acquire copper from Human Serum Albumin (HSA), one of the main Cu(II) carriers in the blood(Jenkitkasemwong et al., 2012; Shenberger et al., 2015; Stefaniak et al., 2018). hCTR1 N-term has several methionine and histidine-rich clusters that are previously shown to be essential for copper acquisition and copper binding(Guo et al., 2004; Larson et al., 2010; Stefaniak et al., 2018). The presence of the **<u>A</u>**mino **<u>T</u>**erminal **<u>Cu</u>**(II)-and **<u>N</u>**i(II) binding site (ATCUN) spanning the first three amino acid residues of hCTR1 and characterized by the general sequence (H_2_N-Xaa-Zaa-His) favors a possibly direct Cu(II) transfer from HSA to hCTR1 (Shenberger et al., 2015; Schwab et al., 2016). Stefaniak *et al*. has indicated that other residues adjacent to the ATCUN site, namely His^5^, His^6^, and Asp^13^ might also be involved in Cu(II) binding along with the ATCUN cluster at a pH characteristic of the extracellular space(Stefaniak et al., 2018). Intracellular copper exists primarily in reduced form, Cu(I), whereas in the extracellular environment, it is found in its higher oxidation state, Cu(II)(Maryon et al., 2007b). It is hypothesized that an extracellular reducing factor such as ascorbate or STEAP reductases might be responsible for the reduction to happen(Öhrvik and Thiele, 2014). Though *in vitro* studies from Haas group have shown that purified N-term of hCTR1 has the capacity to reduce Cu(II) in the presence of ascorbate, any direct role of N-term of hCTR1 in this reduction process and subsequent import is still speculative(Pushie et al., 2015; Galler et al., 2020). This N-terminal extracellular domain contains multiple methionines (M^7^GM^9^, the first Met-cluster and ^40^MMMMPM^45^, the second Met cluster) and histidine (H^3^-H^6^, H^22^-H^24^, and H^31^-H^33^) rich clusters that are possible anchoring sites for Cu(I) and Cu(II) ions respectively. After shuttling through the N-term, the reduced copper is then passed through a Cu(I) specific selectivity filter formed by a conserved ^150^MXXXM^154^ sequence in the second TM domain before it is delivered to cytoplasmic copper chaperone proteins, like Atox1 and CCS(Howell and Abada, 2010; Magistrato et al., 2019).

Using a combination of computational and experimental techniques, we have postulated a model that correlates and links the three main functionalities of the protein, i.e., (a) distinct Cu(I) and Cu(II) binding to hCTR1 N-terminus (b) Cu(I) import and finally (c) hCTR1 endocytosis. We have also highlighted the mechanism by which hCTR1 N-terminal Met clusters in association with the aspartates maintain the reduced redox state of copper, hence rendering it bioavailable without cofactor requirements.

## Results

### hCTR1 facilitates copper uptake at the basolateral membrane in polarized epithelial cells

Due to its unique nature, mammalian polarized epithelial cells connect as well as partition the luminal side (apical) and blood side (basolateral) of the epithelial tissue. Copper is required to be imported in cells from either blood plasma or from luminal contents of the epithelia for its systemic and intracellular utilization.

We used the widely accepted model of polarized epithelia, Madin Darby Canine Kidney (MDCK-II) cells to determine the effect of copper on localization of hCTR1(Gravotta et al., 2019; Perez Bay et al., 2016). To determine the localization of hCTR1 in polarized epithelia, we transfected MDCK cells with Flag-hCTR1, seeded them in tranwell chambers and let it polarize until apical and basolateral domains were formed (Fig S1A, S1B). Cortical actin was used to mark the cell boundary and upon capturing the XZ image (axial plane) of the cells we could clearly distinguish the apical surface from the basolateral surface (Fig S1A). Upon imaging the cells along the axial plane, we found that hCTR1 localizes completely on the basolateral surface with no signal on the apical membrane (Fig1A, *top panel*). Upon copper treatment (100µM; 1hr), hCTR1 endocytoses to vesicular compartments (Fig1A, *bottom panel*). From a range of 0-100µM, optimum copper concentration was determined to be 100uM that triggered complete endocytosis of hCTR1 from plasma membrane to vesicles (Fig.S1C). Using endosomal markers, Rab-GTPases, Rab7 (late), Rab5 (early), Rab9 (endosomes recycling to Golgi) and pulsed transferrin (Tf) uptake assay (5mins and 30 mins), we determined the identity of the endosomal compartments that harbours endocytosed hCTR1. We found that endocytosed hCTR1 primarily localizes at basolateral sorting endosomes (BSE) and the supra-nuclear common recycling endosomes (CRE) compartments marked with Tf post-30 mins uptake (Fig1B).

**Fig.1.**
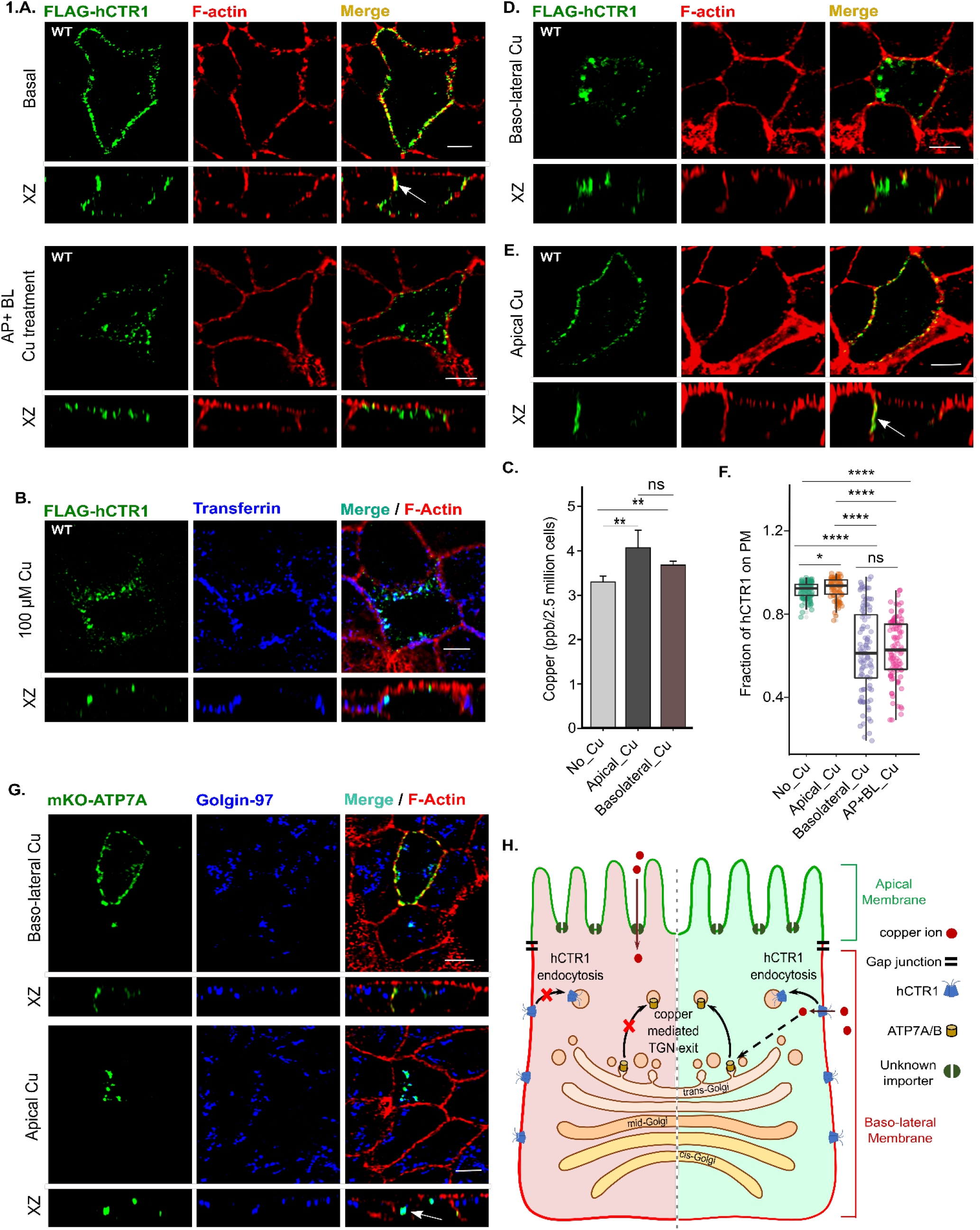
Copper treatment induces hCTR1 endocytosis. (**A)**. Wild-type (WT) FLAG-hCTR1 (green) under basal condition colocalizes with phalloidin (red), white arrow in the XZ section exhibits basolateral localization of hCTR1 (*Upper panel*). WT FLAG-hCTR1 (green) endocytosed upon copper treatment from both apical (AP) and basolateral (BL) sides of the cells (*Lower panel*). **(B)** Following copper (100 μM) mediated endocytosis, WT FLAG-hCTR1 colocalizes with the basolateral sorting endosomes and common recycling endosomes marked with post-30 min internalization of Transferrin-Alexa-633 (Blue). **(C)** Comparison of copper uptake in polarized MDCK-II cells (n=9) under different treatment conditions shows similar copper accumulation in case of both apical and basolateral copper treatment, copper concentrations were measured in parts per billion (ppb, mean ± SD)**P<0.01; ns, not significant (non-parametric Mann–Whitney U test/Wilcoxon rank-sum test) **(D)** WT FLAG-hCTR1 (green) endocytoses in response to copper treatment only on the basolateral side of the cells **(E**) WT FLAG-hCTR1 (green) treated with copper on the apical side fails to endocytose and localizes at basolateral membrane. **(F)** Fraction of hCTR1 colocalization with membrane marker F-actin, demonstrated by a box plot with jitter points. The box represents the 25–75th percentiles, and the median in the middle. The whiskers show the data points within the range of 1.5× interquartile range (IQR) from the 1st and 3rd quartile. *p<0.05, ****P<0.0001; ns, not significant (non-parametric Mann–Whitney U test/Wilcoxon rank-sum test). Sample size (n) for Basal: 100, AP: 91, BL: 101, AP+BL: 101. **(G)** Stably expressed mKO-ATP7A (green) in response to copper treatment at the bottom transwell chamber (*upper panel*) traffics to the basolateral membrane whereas upon copper treatment at the top transwell chamber (*lower panel*) it localizes with Golgin97 (blue). **(H)** Schematic showing the physiological effects of apical versus basolateral copper entry, the latter through hCTR1. [In all the conditions, cells are polarized MDCK-II, XZ section shows the orthogonal sections of all the stacks, green-FLAG-hCTR1 and mKO-ATP7A separately, red-F-actin and Blue-Transferrin and Golgin-97 separately; scale bar-5 μm].

Apart from hCTR1, the divalent metal transporter DMT1 has been implicated as another copper importer, albeit at a low affinity, but at the apical membrane of polarized epithelia(Arredondo et al., 2003). Hence we determined whether uptake of copper happens primarily through the apical or basolateral side of MDCK-II. Cells were polarized on transwells and treated with 100µM copper on either basolateral (bottom chamber) or apical side (top chamber) of the transwells. Using ICP-OES, we measured intracellular copper levels and found that luminal (apical) as well as basolateral membranes can uptake copper with similar efficacies (Fig1C). To determine whether hCTR1 endocytosis is triggered primarily in response to apically or basolaterally imported copper, we treated polarized MDCK cells expressing Flag-hCTR1 either in the top or bottom chamber of the transwells. We observed that upon copper treatment at the bottom chamber, hCTR1 endocytosed (Fig1D), whereas it continued to localize on the basolateral membrane upon top chamber copper treatment (Fig 1E). Quantitation of colocalization between Flag-hCTR1 and F-Actin upon copper treatment at basolateral vs apical sides of MDCK-II is illustrated in Fig1F. To summarize, copper imported through the basolateral membrane triggers hCTR1 endocytosis.

Copper uptake at the basolateral membrane is mediated by hCTR1 and at the apical side by an unknown mechanism or possibly by DMT1(Lee et al., 2002b; Zimnicka et al., 2011). We used CuCl_2_ as the source of copper. Cu(II) needs to be reduced to bioavailable Cu(I) prior to it being utilized by intracellular proteins. We further investigated if both these copper pools (luminal vs basolateral) are equally bioavailable and are able to trigger physiological response inside the cell. We utilized copper-induced trafficking of the Copper-ATPases ATP7A and ATP7B from the trans-Golgi network to secretory vesicles as a readout of bioavailable or utilizable copper. It has been established that upon copper entry via hCTR1, copper is sequestered by metallochaperone, Atox1. Atox1 further transports the copper to Copper-ATPases that traffics to vesicles to export out excess copper(Lutsenko et al., 2007). Upon copper treatment at the basolateral side of MDCK-II we observe TGN exit, vesicularization and plasma membrane targeted trafficking of the ATP7A (Fig1G, *top panel*) as well as ectopically expressed ATP7B (FigS1D, *top panel*). However, copper treatment at the apical side, though lead to intracellular copper uptake, did not elicit any trafficking response of the Copper-ATPases, ATP7A (Fig1G *bottom panels)* or ATP7B (FigS1D *bottom panels*). In conclusion, copper that enters through the basolateral membrane via hCTR1 is bioavailable and therefore can evoke normal physiological response. A schematic summarizing the different physiological responses elicited by apical versus basolateral copper uptake is shown in Fig 1H.

### N-term of hCTR1 is critical for its plasma membrane localization

The N-term of hCTR1 forms the extracellular domain that is exposed to copper from the basolateral side of polarized epithelial cells. Given that, the cytosolic C-term should first sense the apically incorporated copper, however it fails to induce endocytosis of the protein. So we hypothesized that it is the N-term (1-67 amino acids) but not the C-term that primarily senses the extracellular copper and induces subsequent early physiological responses. To test that, we deleted the N-term (1-67) and determined intracellular localization of the truncated protein in basal and copper treated conditions. Interestingly, we found that Δ67-hCTR1 though expresses well and folds properly to exit ER, fails to localize at the basolateral membrane. Instead, Δ67-hCTR1 localizes on the TGN at basal or high copper conditions [Fig 2A (top and bottom panel)]. We can conclude that the N-term is crucial for plasma membrane localization of hCTR1 that is either regulated by copper binding to the N-term or by rendering the protein in TGN-exit favourable conformation. Deleting the entire N-term did not provide us a detailed view of its role in copper sensing and copper uptake. However, previous study by Eisses et al showed that this same truncated form can uptake copper at a very low level in sf9 cells(Eisses and Kaplan, 2005).

**Fig. 2.**
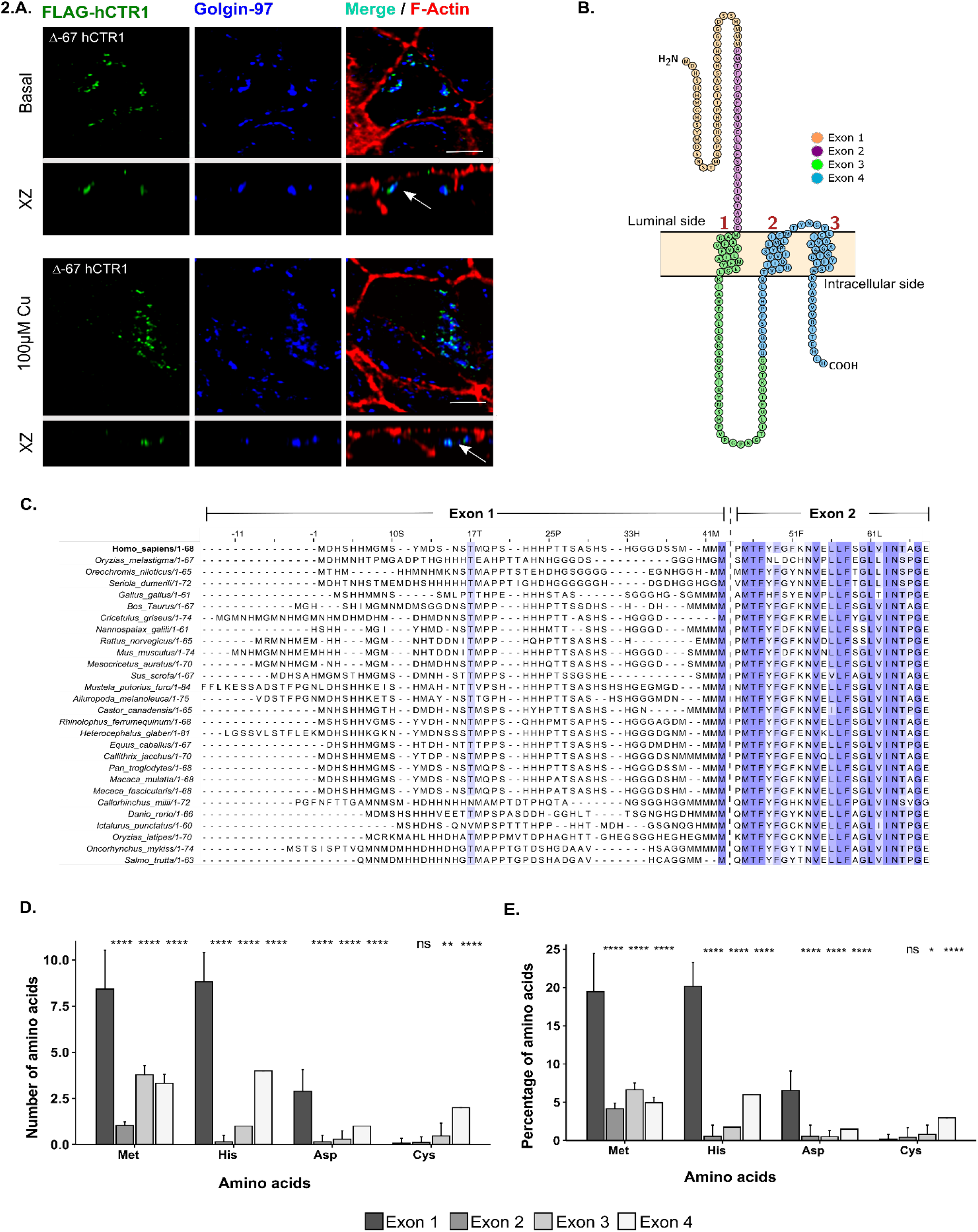
His-Met rich N-term of hCTR1 is crucial for its plasma membrane localization. **(A)** Amino-terminal deleted Flag-Δ-67 hCTR1 (green) colocalizes with TGN-marker Golgin-97 (blue), both in basal (upper panel) and 100μM copper treated condition (lower panel). **(B)** Amino acids of hCTR1 monomer color coded based on the respective exons coding them. Representative image was made in Protter. **(C)** Sequence alignment of the first two exons of chordate CTR1s shows higher variability in the first exon as compared to the second one. Conserved residues are marked by blue. Intensity of blue colour is proportional to the conservation status of a residue. Bar plot (mean ± SD) representing the number **(D)** and percentage **(E)** of His, Met, Asp and Cys residues coded by the four exons of CTR1 from twenty eight chordate species. *p<0.05, **p<0.01, ****P<0.0001; ns, not significant (non-parametric Mann– Whitney U test/Wilcoxon rank-sum test).

### The distal amino-terminal domain exhibits higher phylogenetic variability as compared to the proximal part

A comprehensive sequence alignment of 219 CTR1s across the species and their homology analysis reveals that the extracellular amino-terminal domain is not evolutionarily conserved (FigS2A). In contrast, the hCTR1 TM domains and the cytosolic tail exhibit higher phylogenetic similarity and contain conserved motifs such as the ^150^MXXXM^154^ in TM2 and ^188^HCH^190^ in the cytosolic tail(Guo et al., 2004; Maryon et al., 2013). Chordate CTR1 is mostly composed of four exons, and the domain architecture encoded by the four exons is illustrated in Fig2B. For hCTR1, the first two exons code for the extracellular amino-terminal domain. The third one codes for the first TM (TM1) domain and part of the cytosolic loop flanked by TM1-TM2. The fourth exon encodes for the central pore-forming region of the protein necessary for copper transport and the cytosolic C-terminal domain(Schushan et al., 2010).

A relatively higher conserved amino-acid sequence of the ‘‘channel-pore forming exon” coded by Exons 3 and 4 (illustrated in FigS2B) suggests that the mechanism of copper import is conserved across the species. The higher variability in the first exon could be attributed to the differences in copper sensing and acquiring mechanisms in different species growing in different environments. Interestingly, sequence conservation also varies within the N-term. The distal part (away from the TM1) of the N-term (amino acid 1-42) is less conserved, and the proximal half (towards TM1, from amino acid 43-67) is relatively more conserved among various species (Fig2C). Exon1, despite having higher variability, exhibits bias towards certain residues, namely histidine, methionine, and aspartic acid, in terms of total abundance as well as percentage (Fig2D,2E, FigS2C). As in line with previous studies(Guo et al., 2004), we hypothesize that the higher prevalence of a few amino acids is functionally relevant and participates in copper coordination and transport.

### N-terminal His-Met clusters and aspartate residues are putative Cu(II) and Cu(I) coordination residues

The N-terminal domain of hCTR1 has been predicted to be unstructured(Aller and Unger, 2006; De Feo et al., 2009; Ren et al., 2019). Despite multiple trials using various affinity tags, we failed to purify the full-length N-term of hCTR1 for copper-binding assays. The sequence comparisons presented above suggest that the shorter synthetic peptide (^1^M-^14^S) construct that has been previously reported will provide an incomplete understanding as it lacks the key His-Met motifs (H^22-24^, ^31^HSH^33^, and ^40^MMMMPM^45^) of the proximal N-term(Stefaniak et al., 2018). Consequently, we explore the molecular mechanism of the copper-binding at the N-terminal domain using a combination of classical molecular dynamics and enhanced sampling approaches that provide an accurate computational method to probe the free energy surface of complex processes. In addition, the Cu(II)/Cu(I) ion is represented as a virtual site model that allows us to reproduce the coordination geometry of the metal ion without additional quantum chemical calculations(Liao et al., 2015; Pang, 1999). Several studies have shown the excellent agreement of these enhanced sampling methods with experiments and are thus very well suited to provide critical insights into the binding and unbinding process by virtue of predicting binding constants and molecular mechanisms(Chowdhury et al., 2021). In the first step, we independently simulated copper ion in both oxidation states, Cu(II)/Cu(I), in conjunction with the N-terminal domain (system setup is shown in Fig S3A). Although the ATCUN motif is well known to be responsible for the initial steps of copper-binding, the high structural flexibility and relative abundance of histidine and methionine in spatially nearby regions required a more comprehensive sampling of Cu(I) and Cu(II) association in hCTR1. From the simulations, we observe transient interactions of Cu(II)/Cu(I) with the N-terminal domain. To refine the different interaction sites, well-tempered metadynamics simulations were used to calculate the binding free energy of the Cu(II)/Cu(I) ion along a pathway described by two collective variables (described in Fig S3B and S3C). Details of the simulation are presented in Table S3D.

A schematic elucidating the molecular mechanism of Cu(II) and Cu(I) binding to the N-term as probed through our metadynamics simulations is shown in Fig 3A. The converged free energy profiles of both Cu(II) and Cu(I) consist of a series of minima corresponding to specific sites for ion-protein interactions along the binding/unbinding pathway. The potential binding pathways of the Cu(II) and Cu(I) ion were computed using the nudged elastic band method (Henkelman and Jónsson, 2000), and the main clustered structures were identified (Fig 3B and 3C). Our results indicate that the binding (or equivalently, the unbinding) of the Cu(II) octahedral virtual site model occurs at histidine-rich sites (Fig 3B). The bioinorganic complexes observed contains coordination bonds with imidazole nitrogen atoms from a nearby histidine residue and two carboxylate group from two proximal aspartate residues. The coordination sphere of the octahedral Cu(II) was further complemented by the solvent molecules. The subsequent unbinding of copper gives rise to a highly hydrated Cu(II) state commonly bound to a single carboxylate group only. For Cu(II), log K has been calculated to be 13.66 from the most probable binding pathway on the computed and converged free energy surface. Considering K_d_ = 1/K, the dissociation constant, when converted to free energy at 300K temperature, yields a value of 18.76 kcal/mol. Our data matches quite well with that of Stefaniak *et al.,* who has provided a log K value of 13.2 ±0.3, for Cu(II) binding to the model peptide hCTR1_1–14_ through NTA competition assay(Stefaniak et al., 2018); on the contrary, a much lower log K value for Cu(II)-hCTR1_1–14_ as well as for Cu(II)-hCTR1_1–55_ have also been reported in the literature, which might not be exact, given that they did not explicitly take into account the interference of buffer and other solution components(Haas et al., 2011; Du et al., 2013).

**Fig 3:**
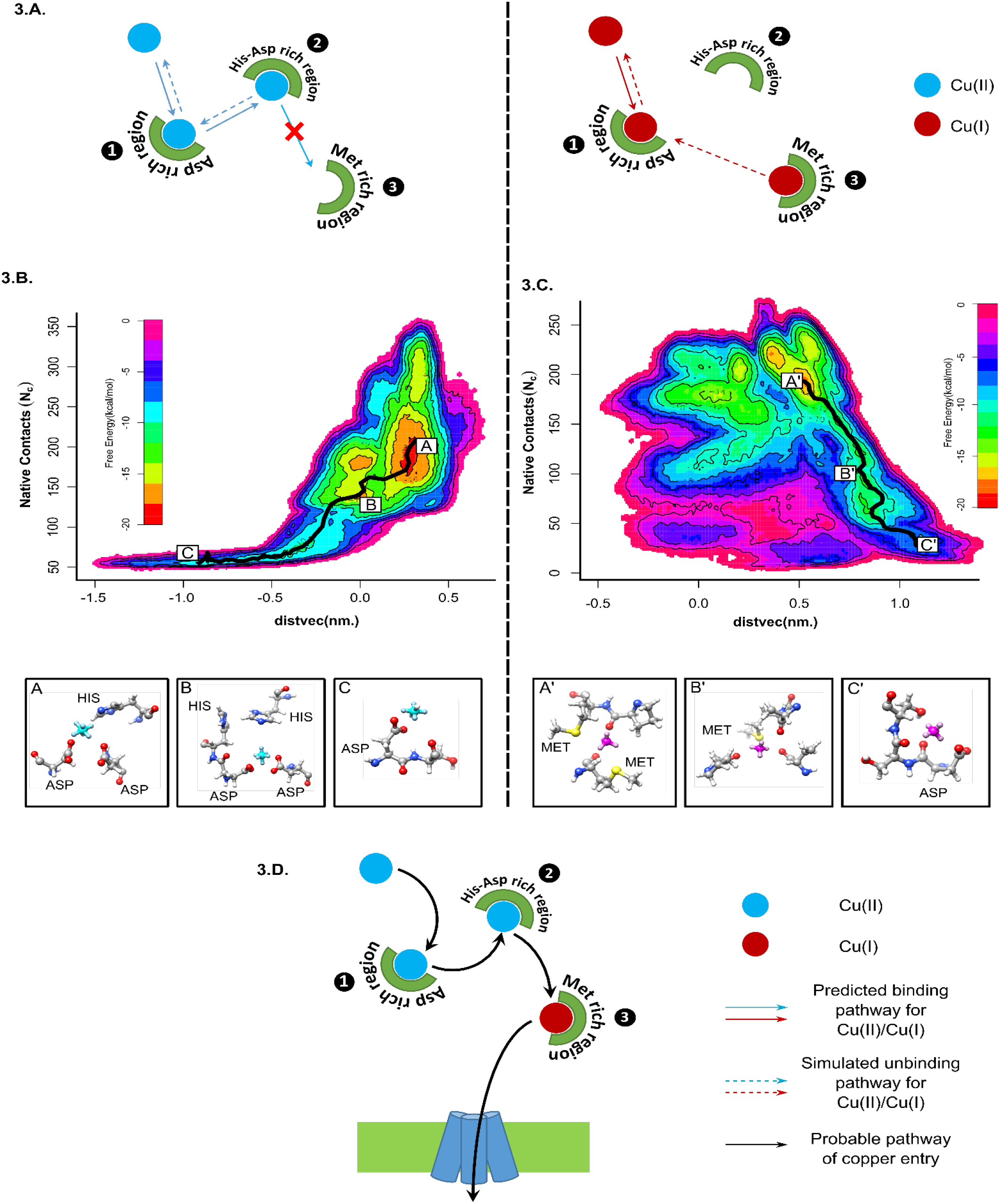
N-term aspartic acids and histidines coordinate Cu(II) whereas methionines are primarily responsible for binding to Cu(I). **(A)** Schematic of the molecular mechanism as probed through metadynamics simulations for both Cu(II) (left panel) and Cu(I) (right panel). Free energy surface of Cu(II) **(B)** and Cu(I) **(C)** binding to trimeric hCTR1 N-term (67 amino acids) against two collective variables, Native contacts (*NN*_C_) and distvec (nm). The structures along the path of dissociation are shown below the free energy diagram (A, B and C indicates the different Cu(II) coordination complexes whereas A’, B’ and C’ indicates the different Cu(I) complexes). Cu(II) octahedral model is shown in cyan and the Cu(I) tetrahedral model is shown in magenta. **(D)** Schematic of the proposed pathway of copper shuttling through the N-term.

In the next step, we performed a similar metadynamics simulation for the tetrahedral Cu(I) virtual site model to analyze the binding of the Cu(I) ion at the N-terminal domain. Upon inspection of clustered structures on the free energy surface along the minimum free energy path (Fig 3C), we observe that Cu(I) binding occurs in exclusively methionine-rich regions, in stark contrast to the computationally predicted complexes of Cu(II). The bound-state contains up to two methionine residues coordinated *via* their sulfur atoms. Upon unbinding, the complexes transition into relatively oxygen donor rich complexes before becoming fully hydrated and thus fully unbound. In the case of Cu(I), our calculated log K comes around 13.17, which at 300 K corresponds to a free energy value of 18.084 kcal/mol. However, our data is not in very good agreement with studies carried out by Du *et al*. and Yang *et al*., where they observe log K values of 14.92 and 14.7 for Cu(I) for hCTR1_1–55_ and hCTR1_1–46_ respectively through competition reactions with bicinchoninic acid (BCA)(Du et al., 2013; Yang et al., 2019). The difference in the values of dissociation (or binding) constants might arises due to the difference in the reference considered. The free energy from our simulations is calculated considering a completely solvated structure of Cu(I) as the fully unbound state, whereas under experimental conditions, Cu(I) always remains in complex with some ligands and never as just a fully hydrated ion.

Our results suggest that both Cu(II) and Cu(I) can bind to the N-terminal domain with very disparate binding modes. This is also the first time we show that there is a possible role of aspartate residues found abundantly on hCTR1 N-term, primarily in the Cu(II) binding process. This hypothesis is further tested by experiments in the following sections. A probable pathway of Cu(II) and Cu(I) shuttling through the N-term before it enters the TM pore is shown in Fig 3D.

### N-terminal methionine cluster and aspartate facilitates copper uptake

To understand the role of copper binding to the amino-terminal residues (predicted in the previous section) in its subsequent uptake by hCTR1, we utilized yeast complementation assay. We expressed the WT hCTR1 and multiple hCTR1 N-terminal mutants (illustrated in Fig 4A) in ΔyCTR1 (*BY4742 Saccharomyces cerevisiae* strain lacking endogenous yCTR1). Yeast growth rate phenotype was used as an indicator of copper uptake in restrictive media(Wu et al., 2009). We measured yeast growth in non-fermenting sugar (ethanol and glycerol) containing media as it will allow the growth of only those colonies which are able to import copper through ectopically expressed hCTR1 (WT or mutant hCTR1). Subsequently, that imported copper will activate the cytochrome c oxidase of mitochondria for ATP production. Thus, cell growth in this YPEG (Yeast extract, Peptone, Ethanol, and Glycerol) media is proportional to the copper import property of the transformed hCTR1 constructs (details in Materials and Method). We measured yeast growth both qualitatively and quantitatively (in the plate and in liquid culture media, respectively). Empty pTEF vector transformed strain showed no growth in YPEG plate, and WT hCTR1 (cloned in pTEF) recovered the growth indicates that human CTR1 is able to complement yCTR1 (Fig4B). There are three separate clusters of histidines, ^3^HSHH^6^, H^22-24^, ^31^HSH^33,^ that are postulated to bind Cu (II). In solid media culture using colony counting, we found that ΔH^3^-H^6^ and ΔH^22^-H^24^ mutants yeast exhibited similar growth patterns to the WT hCTR1 (data not shown for ΔH^22^-H^24^). Lower colony counts in ΔM^7^-M^9^ and D13A mutants indicate reduced growth in comparison to WT (Fig4B). Growth kinetics was measured during the log phase in media culture that is otherwise not possible in end-point colony counting in plate cultures. Replicating the data from colony counts, ΔM^7^-M^9^ and D13A showed a reduced growth rate, i.e., 36% and ∼15%, respectively, than the WT indicating reduced copper import. ΔH^3^-H^6^ expressing yeast showed no alteration in the growth rate compared to the one expressing WT-hCTR1, which signifies that copper uptake of this mutant is unaltered. ΔM^40^-M^45^ mutant showed ∼15% reduced growth as compared to the WT (Fig4C). To summarize, the methionine clusters, especially the first one, are the key motif that participates in copper uptake.

**Fig 4.**
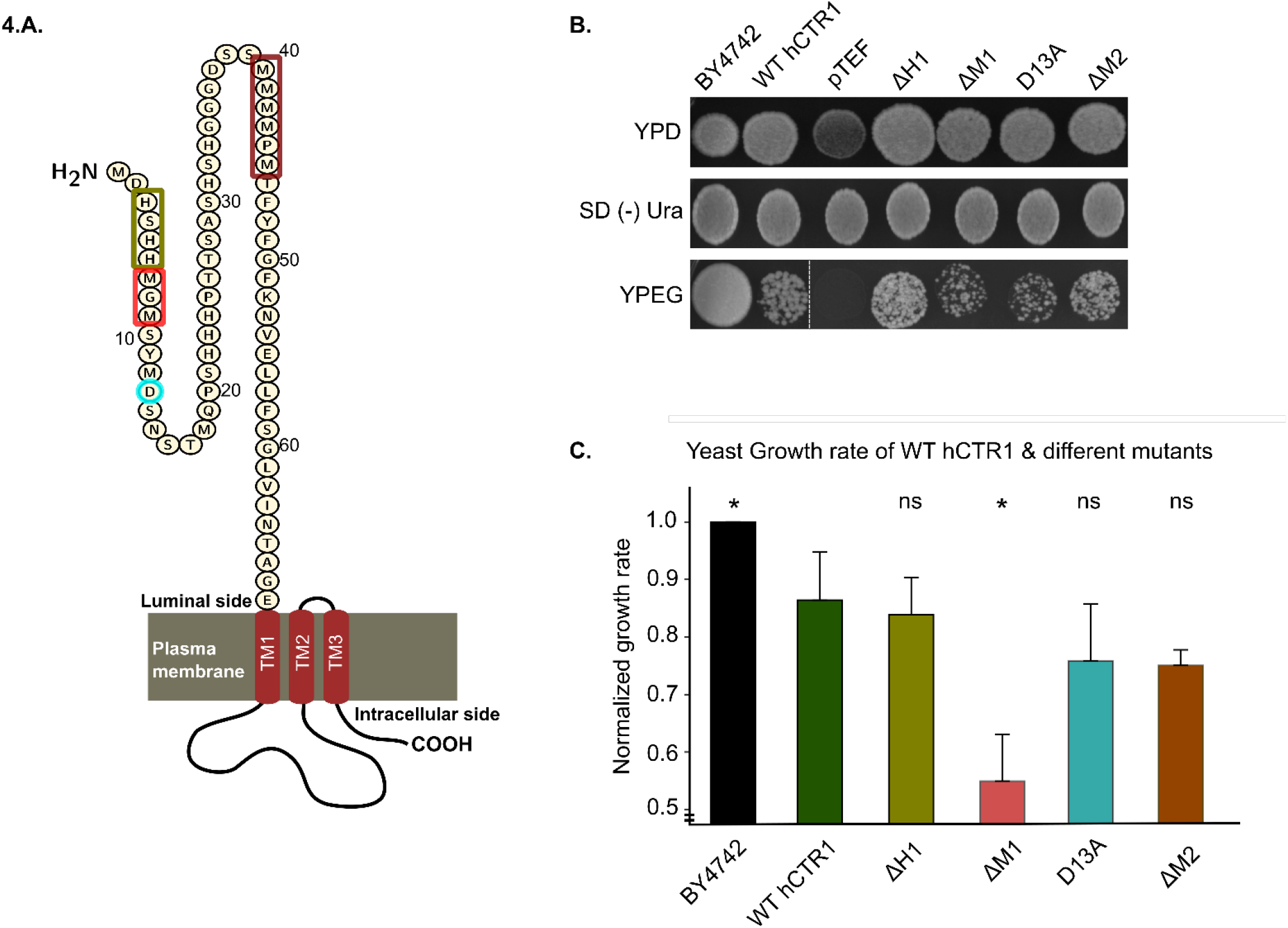
hCTR1 N-term methionine and aspartic acid residues are critical for copper import. **(A)** Illustration of the 67 amino acid long hCTR1 N-term; colour-coded rectangles and circles indicate the position of the mutations used in yeast complementation assays. **(B)** In the YPD plate, all the constructs show appreciable growth (*top panel*). Mutants selected in SD (-) Ura plate following transformation. In the YPEG plate, the empty vector (pTEF) containing strain (ΔyCTR1BY4742) fails to grow but WT-hCTR1 is able to successfully complement the ΔyCTR1 growth defect. **(C)** Normalized growth rate of WT and mutant hCTR1s (mean ± SD) shows significantly reduced growth in ΔM1 (ΔM7-M9) mutant, followed by D13A and ΔM2 (ΔM40-M45). ΔH1 (ΔH3-H6) shows almost similar growth rate as that of WT. *p<0.05, ns, not significant (non-parametric Mann–Whitney U test/Wilcoxon rank-sum test w.r.t WT hCTR1). Sample size for each set, n=4. BY4742 is the wild-type yeast strain, containing endogenous yCTR1 and in the remaining cases wild-type and mutant hCTR1s are separately transformed into the yCTR1 deleted strain.

### 7MGM9 and D^13^ are indispensable for copper-induced hCTR1 endocytosis

As a self-regulatory mechanism to limit intracellular copper concentration, hCTR1 endocytoses from the plasma membrane upon copper treatment(Petris et al., 2003). We investigated if copper binding to the N-term, copper import and endocytosis are linked and interdependent to maintain the proper functioning of the protein. To study that, at the outset, we deleted the first His motif, ^3^HSHH^6^ (ΔH1). In agreement with unaltered copper uptake observed using yeast complementation assay, the mutant localizes on the basolateral membrane in basal condition and endocytoses in elevated copper (100μM) comparable to the wild type (WT) protein (Fig 5A). Interestingly, even upon deletion of both the His cluster ^31^HSH^33^and ^3^HSHH^6^ (double mutant, ΔH1H2), we did not notice any deviation in phenotype in basal or elevated copper compared to the wt-CTR1 and ΔH^3^-H^6^ (Fig 5B). Based on our MD findings, we hypothesize that the His motifs sequesters Cu(II) from the environment and increases its local availability for further reduction and uptake. So the role of His motifs would be more apparent in copper limiting conditions. Upon treating the cells with lower copper (25μM), both the above-mentioned single His motif mutant as well as the double His motif mutant failed to endocytose though WT protein endocytoses under similar low copper concentrations (Fig 5C). A similar observation was made in the 5µM copper treatment (data not shown). These results point to an essential role of the N-terminal histidine motifs in acquiring copper under physiological low copper concentrations and thereby promoting hCTR1 endocytosis. Presence of high copper, i.e., 100µM, ‘undermines’ the importance of histidine motifs as the Met clusters can as well bind and sequester Cu(II), though with a lower affinity. Quantitation of endocytosis of the WT and the two His mutants under different copper concentrations has been illustrated in Fig 5D. A schematic summarizing the phenotypes of the WT and the mutants under low and high copper conditions has been elucidated in Fig 5E.

**Fig 5:**
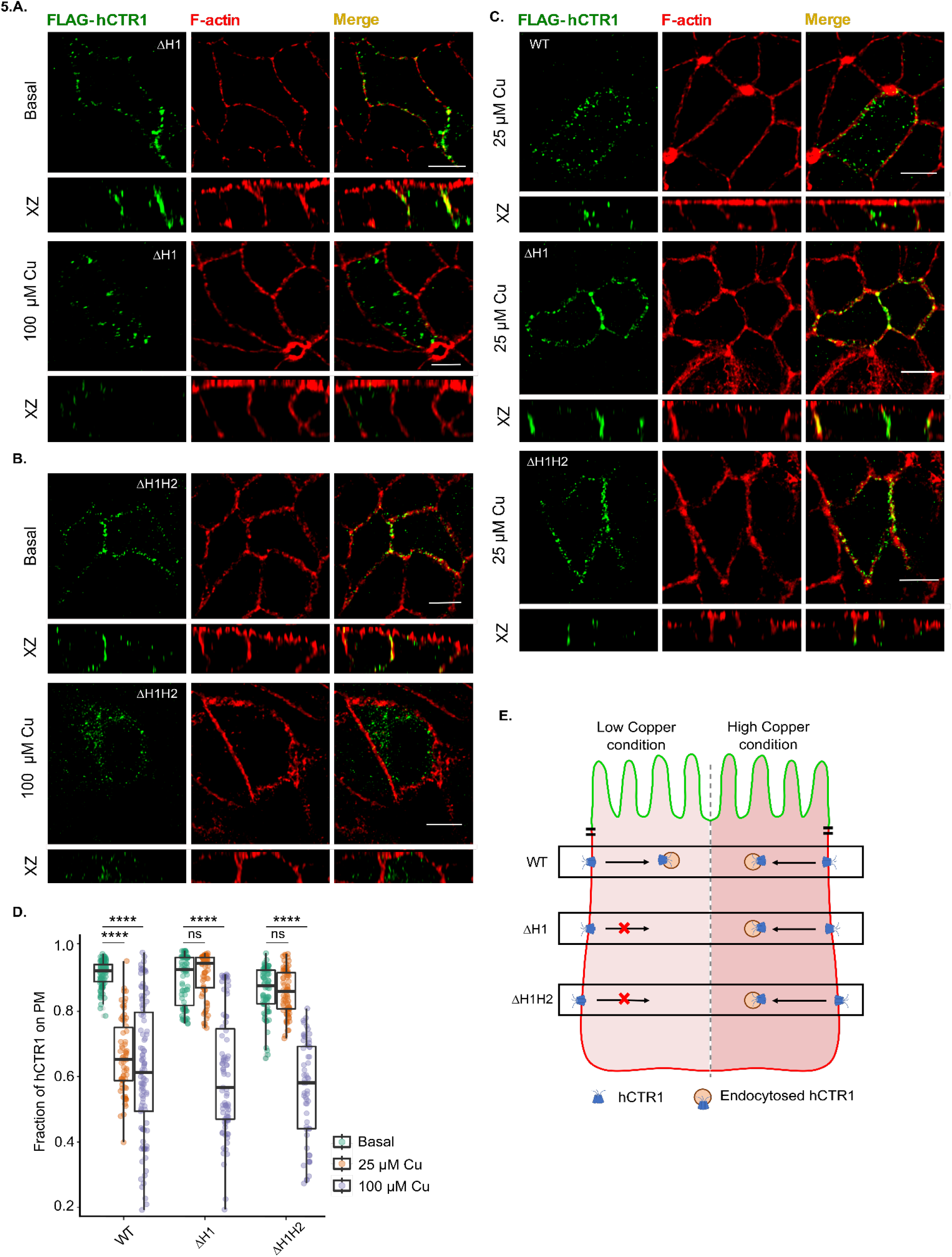
N-term histidines are crucial for hCTR1 endocytosis under copper-limiting conditions. **(A)** ΔH1 Flag-hCTR1 (green) localizes at the basolateral membrane at basal copper (*upper panel*) and endocytoses upon 100 μM copper treatment (*lower panel*). **(B)** ΔH1H2 (combined deletion of ΔH3-H6 and ΔH31-H33) under basal (no copper treatment) condition resides on the PM (upper panel). During 100μM copper treatment, this mutant shows endocytosis (lower panel). **(C)** WT hCTR1 (upper panel) endocytoses under low (25μM) copper treatment, whereas, under similar treatment conditions, both ΔH1 (middle panel) and ΔH1H2 (lower panel) mutants fail to do so. **(D)** Fraction of hCTR1 colocalization with membrane marker F-actin, demonstrated by box plot with jitter points under basal (green circles), 25μM copper (orange circles) and 100μM copper treated (purple circles) conditions. The box represents the 25–75th percentiles, and the median in the middle. The whiskers show the data points within the range of 1.5× interquartile range (IQR) from the 1st and 3rd quartile. ****P<0.0001; ns, not significant (non-parametric Mann–Whitney U test/Wilcoxon rank-sum test). Sample size (n) for WT (basal: 100, 25μM Cu: 66, 100μM Cu: 101), ΔH3-H6 (basal: 78, 25μM Cu: 70, 100μM Cu: 83), ΔH1H2 (basal: 76, 25μM Cu: 77, 100μM Cu: 56). [In all the conditions, cells are polarized MDCK-II, XZ section shows the orthogonal sections of all the stacks, green-FLAG-hCTR1 and red-F-actin; copper treatment-25μM and 100μM separately on the basolateral chamber of the transwell; scale bar-5 μm]. **(E)** Schematic summarizing the phenotypes of the WT and the His-mutants under low and high copper conditions.

hCTR1 contains two N-terminal methionine clusters (M1; ^7^MGM^9^) and (M2; ^40^MMMMPM^45^). To determine their role in the localization and copper-induced endocytosis of the protein, we deleted M1 and M2 individually (ΔM1 and ΔM2) and both the clusters together (ΔM1M2). In basal copper conditions, all these three mutants localized on the plasma membrane. Interestingly upon copper treatment, ΔM1 failed to endocytose from the basolateral membrane (Fig 6A). Moreover, raising the copper concentration to 200µM did not correct the phenotype (data not shown). A similar non-endocytosing phenotype was also observed in the double mutant ΔM1M2 (Fig 6B). Interestingly, upon deleting M2 only, the protein endocytosed upon copper treatment, similar to WT-hCTR1 (Fig 6C).

**Fig. 6.**
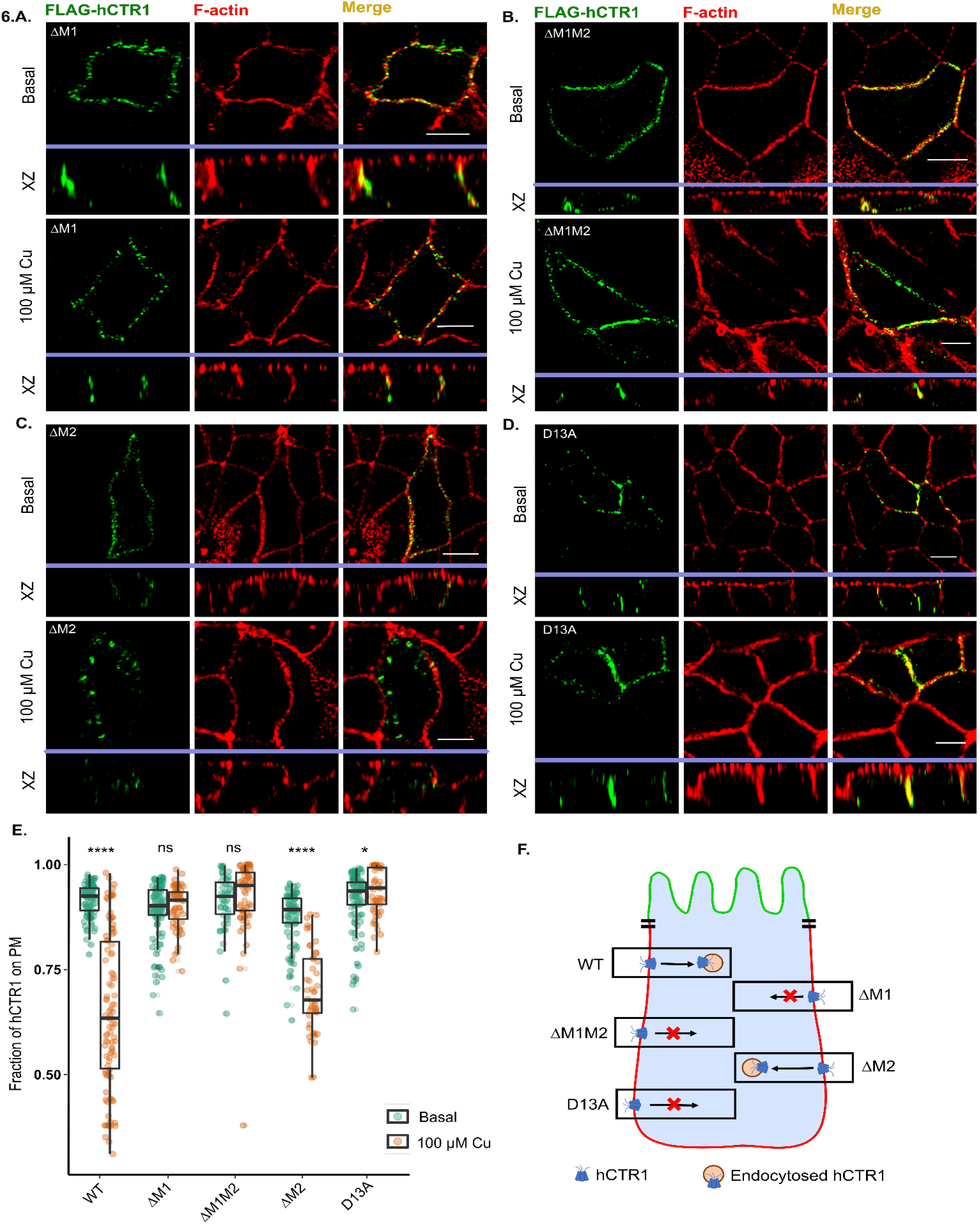
hCtr1 N-term methionines and aspartic acid regulates endocytosis of hCTR1. **(A)** ΔM1 Flag-hCTR1) localizes at the basolateral membrane at basal copper (*upper panel*), and fails to endocytose under treatment with 100 μM copper (*lower panel*). **(B)** ΔM1M2 Flag-hCTR1 (combined deletion of ΔM7-M9 and ΔM40-M45) localizes at the basolateral membrane at basal copper (*upper panel*), it fails to endocytose under treatment with 100 μM copper (*lower panel*). **(C)** ΔM2 Flag-hCTR1 (ΔM40-M45) localizes at the basolateral membrane at basal copper (*upper panel*) and endocytoses upon 100 μM copper treatment (*lower panel*). **(D)** D13A localizes at the basolateral membrane at basal copper (*upper panel*) and fails to endocytose upon 100 μM copper treatment (*lower panel*). (**E)** Fraction of hCTR1 colocalization with membrane marker F-actin, demonstrated by box plot with jitter points under both basal (green circles) and 100 μM copper treated (orange circles) conditions. The box represents the 25–75th percentiles, and the median in the middle. The whiskers show the data points within the range of 1.5× interquartile range (IQR) from the 1st and 3rd quartile. *p<0.05, ****P<0.0001; ns, not significant (non-parametric Mann–Whitney U test/Wilcoxon rank-sum test). Sample size (n) for WT (basal: 100, Cu: 101), ΔM1 (basal: 109, Cu: 61), ΔM1M2 (basal: 51, Cu: 65), ΔM2 (basal: 93, Cu: 48), D13A (basal: 106, Cu: 57). [In all the conditions, cells are polarized MDCK-II, XZ section shows the orthogonal sections of all the stacks, green-FLAG-hCTR1 and red-F-actin; 100 μM Cu treatment on the basolateral (bottom) chamber of the transwell, scale bar-5 μm]. **(F)** Schematic summarizing the phenotypes of the WT and the different Met and Asp-mutants under high copper conditions.

Our simulation studies, for the first time, also indicated the importance of aspartates explicitly for Cu(II) coordination. The N-term contains three aspartate residues, namely-Asp^2^, Asp^13^, and Asp^37^. Asp^2^ is considered to be part of the well-known ATCUN-cluster (comprising the first three amino acids, Met-Asp-His) which is thought to be indispensable for Cu(II) binding(Gonzalez et al., 2018). On substituting Asp^13^ by Ala (D13A), we found that the mutant failed to endocytose under elevated copper conditions (Fig 6D). Quantitation of endocytosis of the Met and the Asp-mutants under basal and high copper treatment conditions has been illustrated in Fig 6E. A schematic elucidating the phenotypes of the WT and the above-mentioned mutants under copper excess conditions has illustrated shown in Fig 6F.

### Cu(I) and not Cu(II) triggers hCTR1 endocytosis

We know that intracellular copper remains bound to proteins in its reduced form (+1), whereas in the extracellular milieu, copper mostly exists in its higher oxidation state (+2)(Wezynfeld et al., 2019; Kirsipuu et al., 2020). Serum albumin has been demonstrated to bind Cu(II) and deliver it directly to the N-term of hCTR1(Wezynfeld et al., 2019). We speculated a role of the hCTR1 N-term, in regulating the reduction of Cu(II) to Cu(I), along with external reducing agents like ascorbate or STEAP(Lee et al., 2002a; Ohgami et al., 2006). Ascorbate has been traditionally used to reduce Cu(II) to Cu(I). However, prolonged and inadvertent exposure to oxidizing agents during an experimental procedure can re-oxidize Cu(I) to Cu(II) even in the presence of ascorbate. Additionally, uncoordinated Cu(I) exhibits very low solubility in aqueous media.

Subsequently, to distinguish between residues involved in differential Cu(II) and Cu(I) binding, we decided to use THPTA (3 [tris(3-hydroxypropyltriazolylmethyl)amine), a water-soluble, highly effective ligand for copper-catalyzed azide-alkyne cycloadditions (CuAAC), along with CuCl_2_ and ascorbate to provide the cells with a direct source of Cu(I). THPTA not only helps in maintaining the Cu(I) oxidation state during the experimental tenure against aerial oxidation, it also enables the protection of biomolecules from oxidative damage by ROS species that might be generated during the reaction(Hong et al., 2009, 2010). UV-Vis spectroscopy of CuCl_2_ along with THPTA (2.0 equivalence) and ascorbate (varying from 0.5-2.0 equivalence) showed that copper was reduced by the entire range of ascorbate used (Fig S4A). Copper remains in its reduced form during the full duration of the treatment (30 mins) with no trace of Cu(II), leading us to hypothesize that it is Cu(I) and not Cu(II) that actually promotes the endocytosis of this copper transporter. Reoxidation of Cu(I) by H_2_O_2_ at the end of 30 mins leads to the reappearance of the Cu(II) peak. (Fig S4B). Cu(II) gives an absorption maximum at 800 nm as evident from the UV-Vis spectrum, which remains intact after the addition of 2 equivalence of THPTA, whereas further addition of 2 equivalence of ascorbate to the previous mixture causes the peak to vanish, indicating that Cu(I) has formed (Fig S4C). Electronic Paramagnetic Resonance (EPR) data also confirmed our UV-Vis experiments (Fig 7A). Taking into consideration that the so-formed Cu(I)-THPTA complex can donate Cu(I) to hCTR1 N-term, we treated the polarized MDCK-II cells expressing Flag-WT-hCTR1 with the mixture of ascorbate: THPTA: CuCl2 in 2:2:1 ratio in HBSS for 30 mins. Conforming to our hypothesis, hCTR1 was found to endocytose in response to this Cu(I) treatment (Fig 7B). We can summarize that, under physiological conditions, when provided by a source of Cu(II), the protein in combination with other external reducing agents reduces Cu(II) to Cu(I), and Cu(I) acts as the main inducer of hCTR1 endocytosis.

**Fig. 7.**
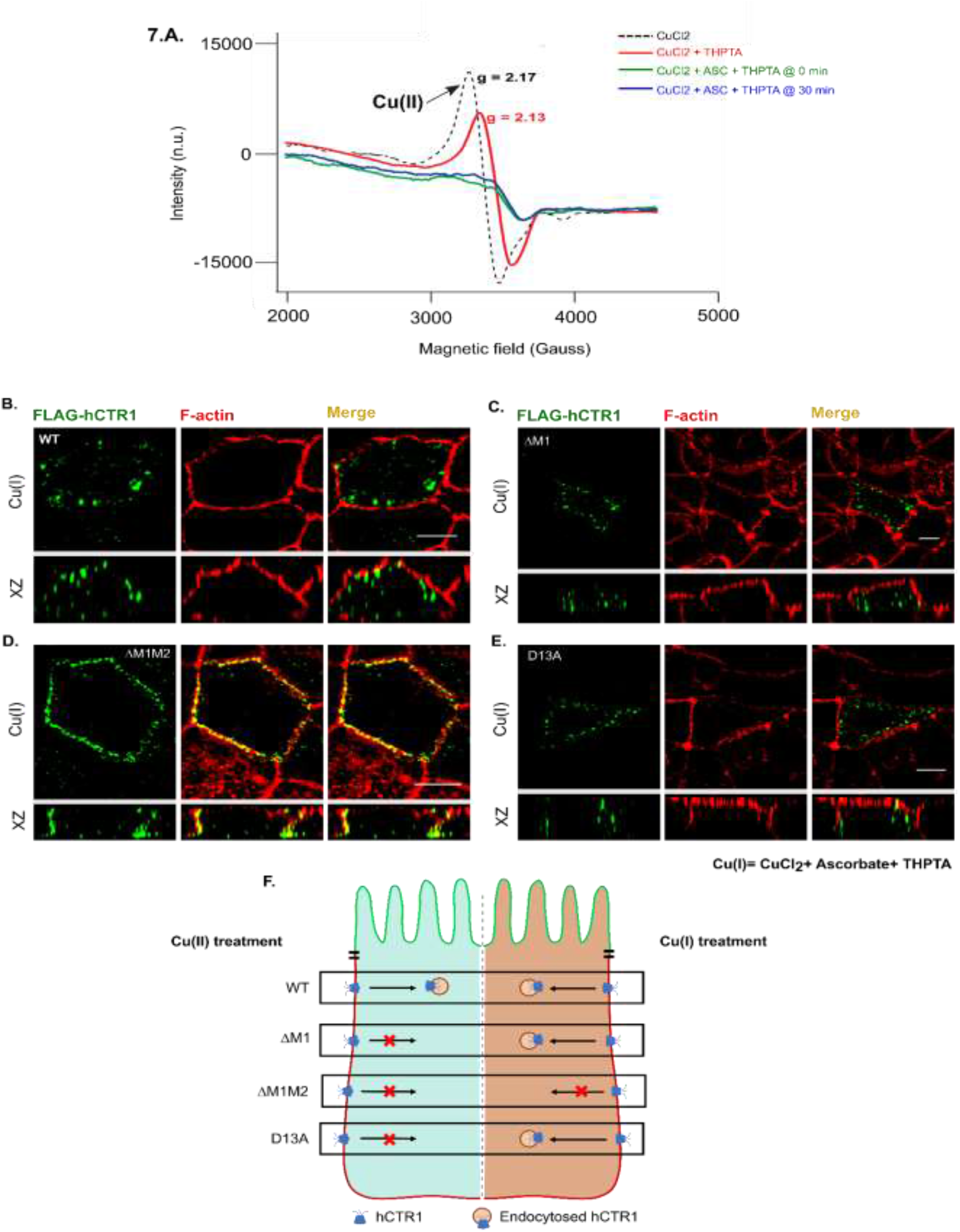
Cu(I) treatment triggers endocytosis of the otherwise non-endocytosing hCTR1 amino-terminal mutants. **(A)** EPR data shows the absence of Cu(II) and presence of only Cu(I) under ascorbate +THPTA + CuCl_2_ (2:2:1) treatment. The peak for Cu(II) does not reappear even after 30 mins which is the entire duration of our following in-vivo experiments **(B)** WT hCTR1 (green) endocytoses when treated with ascorbate +THPTA + CuCl_2_. **(C)** Non-endocytic ΔM1 mutant endocytoses when treated with Cu(I) **(D)** ΔM1M2 Flag-hCTR1 retains its non-endocytic phenotype even under Cu(I) treatment (as marked by its colocalization with phalloidin staining F-Actin (red) (**E)** Non-endocytic D13A Flag-hCTR1 mutant endocytoses when treated with Cu(I). [In all the conditions, cells are polarized MDCK-II, XZ section shows the orthogonal sections of all the stacks, green-FLAG-hCTR1 and red-F-actin; (200 μM ascorbate + 200 μM THPTA + 100 μM CuCl_2_) treatment on the baso-lateral chamber of the transwell, scale bar-5 μm]. **(F)** Schematic summarizing the phenotypes of the WT and the different met and asp-mutants under Cu(II) and Cu(I) treatment conditions.

### Cu(I) treatment induces endocytosis of the otherwise non-endocyting ΔM1 and D13A mutant

As determined in the last section, hCTR1 mutants ΔM1 and ΔM1M2 failed to endocytose under elevated copper conditions (Cu(II) treatment). Based on our MD prediction that methionine clusters bind to Cu(I), we hypothesize that these clusters might play a role in maintaining the Cu(I)-Cu(II) redox balance. We treated the methionine mutants ΔM1 and ΔM1M2 with the mixture of ascorbate: THPTA: copper in a 2:2:1 ratio which acts as a direct source of Cu(I) (as mentioned earlier). Interestingly, ΔM1 is endocytosed (Fig 7C), whereas the ΔM1M2-hCTR1 stays localized at the basolateral membrane (Fig 7D). Upon endocytosis in response to Cu(I) treatment, ΔM1 behaves similar to the WT-hCTR1 and localizes in the CRE as determined by Transferrin uptake assay (Fig S4D). It can be inferred that the presence of at least one methionine cluster is required to bind to the Cu(I) species for its uptake and subsequent endocytosis of the protein.

D13A hCTR1, which remained localized on the PM under Cu(II) treatment, endocytosed in response to Cu(I) (Fig 7E). This observation reinforces our finding that just like the Met cluster, the presence of at least one aspartate residue is required to promote hCTR1 endocytosis. We hypothesize that the aspartates might act as a transient binder for copper in both of its oxidation states during the reduction process and thereby facilitates shuttling of the ion from its histidine-bound +2 state to the adjacent methionine-bound +1 oxidation state. A schematic summarizing the contrasting phenotypes elucidated by the different Met and Asp-mutants under Cu(II) and Cu(I) treatment is shown in Fig 7F.

### His-Met-Asp motifs of the proximal and distal N-terminal domain functionally complement each other

Analysis of the disposition of copper-coordinating amino acids on the N-term and subsequent experimental evidence point towards a possible functional complementarity of region 1-30 and 31-67 residue in the N-term of hCTR1. A previous study by the Thiele group has shown that following copper-induced endocytosis and a cathepsin B mediated truncation of hCTR1 occurs at the amino terminus(Öhrvik et al., 2016; Maryon et al., 2009). Maryon *et al*. showed that hCTR1 undergoes O-linked glycosylation at Thr^27^, and blocking it facilitates a cleavage between the residues A^29^ and G^34^, giving rise to a truncated protein (lacking ∼1^st^ 29 amino acids) that endocytoses to Rab9 compartments in response to high copper(Maryon et al., 2009). Though the truncated protein recycles back to the plasma membrane, it shows reduced copper-import property(Eisses and Kaplan, 2005). To study the individual contribution of the proximal and the distal part of the N-term in copper transport leading to hCTR1 endocytosis, we generated the truncated construct, i.e., Δ30-hCTR1. In basal copper, the deletion mutant localized to the basolateral membrane and endocytosed at elevated Cu(II) and Cu(I) conditions, mimicking the WT-hCTR1 phenotypes (Fig8A, FigS5A). Since in the Δ30-hCTR1 protein, M^7^-M^9^ (M1) is absent, we hypothesize that the reduction of copper, as well as Cu(I) binding, is facilitated by the second methionine stretch, ^40^MMMMPM^45^ (M2). Whether M2 can participate in the reduction or not probably depends on its extracellular solvent accessibility. The presence of the first 30 amino acids possibly causes a steric hindrance affecting the solvent accessibility of the proximal 31-69 amino acid stretch (closer to the transmembrane domain). Hence ΔM1-hCTR1 lacking only M^7-9^ but retaining the rest of the amino acids of the amino-terminal distal half fails to facilitate reduction of Cu(II) and thereby fails to endocytose (Fig 6A). However, in Δ30-hCTR1, the lack of the distal 1-29 residue stretch provides easy access of the proximal ^40^MMMMPM^45^ (M2) to copper in the culture media. In summary, M2 can functionally complement M1, though M1 is considered the principal mediator of endocytosis for the WT protein. In that line of thought, we generated Δ30hCTR1-ΔM2 that lacks both the Met clusters. Similar to the Met double mutant (ΔM1M2), Δ30hCTR1-ΔM2 failed to endocytose both under Cu(II) and Cu(I) conditions, supporting our hypothesis that indeed the two Met-clusters complement each other and at least one Met cluster ensures ‘near proper functioning of the protein (Fig 8B).

**Fig. 8.**
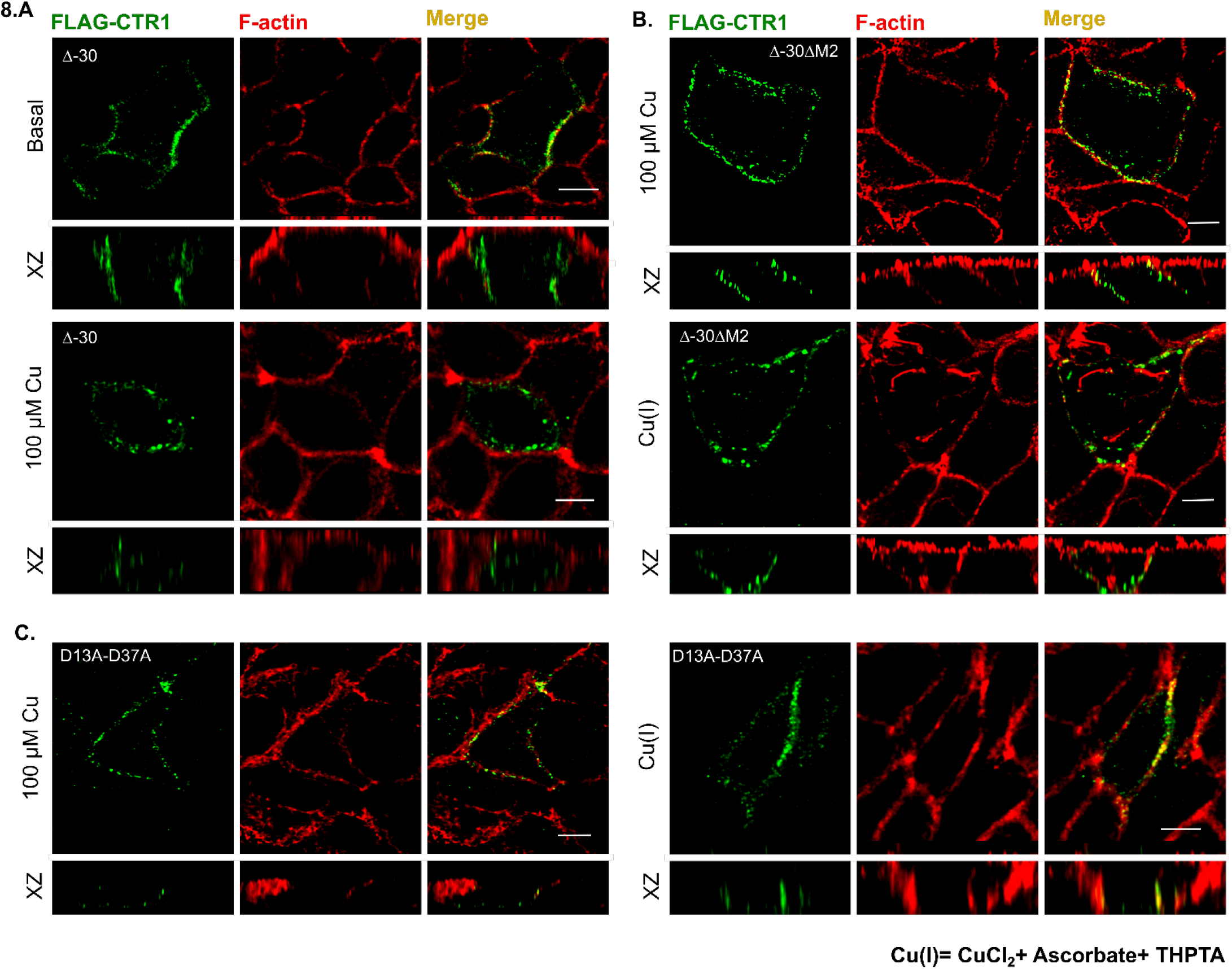
Methionine clusters and Aspartate residues on the proximal and distal part of the hCTR1 amino-terminal exhibit complementarity. (**A)** Δ-30-Flag-hCTR1 localizes at the basolateral membrane at basal copper (*upper panel*) and endocytoses in high copper (*lower panel*) (**B)** Δ30-ΔM2 Flag-hCTR1 mutant failed to endocytose when treated with both Cu(II) (*upper panel*) as well as with Cu(I) (*lower panel*), reminiscent of the phenotype exhibited by ΔM1M2 (*see* Fig 6B *and* 7D *respectively*); **(C)** D13A-D37A Flag-hCTR1 mutant failed to endocytose when treated with both Cu(II) (*left panel*) as well as with Cu(I) (*right panel*). [In all the conditions, cells are polarized MDCK-II, XZ section shows the orthogonal sections of all the stacks, green-FLAG-CTR1 and red-F-actin; 100 μM Cu and (200 μM ascorbate + 200 μM THPTA + 100 μM CuCl_2_) treatment on the baso-lateral chamber of the transwell, scale bar-5 μm].

In our previous experiment with 100μM copper, Δ^3^HSHH^6^ showed copper-induced endocytosis (Fig 5A). We investigated whether deleting the His cluster present on the proximal part of the N-term shows a similar phenotype. We observed that Δ30hCTR1-Δ^31^HSH^33^ (Δ30hCTR1-ΔH2) mutant exhibits normal localization on the plasma membrane and subsequent endocytosis upon 100 μM copper treatment similar to the WT protein and the Δ^3^HSHH^6^ mutant (Fig S5B, *top and bottom panel*). Under low copper conditions (25μM), however, this mutant failed to endocytose completely and remained localized on the PM ((Fig S5B, *middle panel*).

As mentioned in the previous section, the single Asp mutant (D13A) failed to endocytose under Cu(II) treatment, whereas under direct Cu(I) application, it showed WT phenotype. Mutant 2D-2A (D13A-D37A), lacking both D13 and D37, failed to endocytose under both Cu(II) as well as Cu(I) treatment (Fig 8C, left and right panels, respectively). This observation is in agreement with our previous hypothesis that the aspartates (D13 and D37) indeed play a crucial role in transiently binding and shuttling of Cu(I) to the nearest Met clusters (M1 and M2), respectively. Therefore, we can conclude that in the D13A mutant, D37 functionally complements D13 along with the adjacent methionine stretches, M^40^-M^45,^ in respectively transferring copper and binding Cu(I), thereby facilitating its uptake. The presence of at least one of either aspartates, D13 or D37, is warranted for the uptake of copper.

To summarize, the His-Met clusters and the Asp residues on the distal part and the proximal part of the N-term exhibits complementarity. In the absence of the distal amino acid stretch harbouring the first His-Met-Asp cluster, the second cluster can possibly perform the function of the first one in maintaining the redox state of copper that facilitates its uptake and subsequent endocytosis of hCTR1. Our findings provide an explanation of how the truncated version of hCTR1 that lacks O-linked glycosylation maintains functionality as a copper transporting-recycling protein (illustrated in Fig 9).

**Fig. 9.**
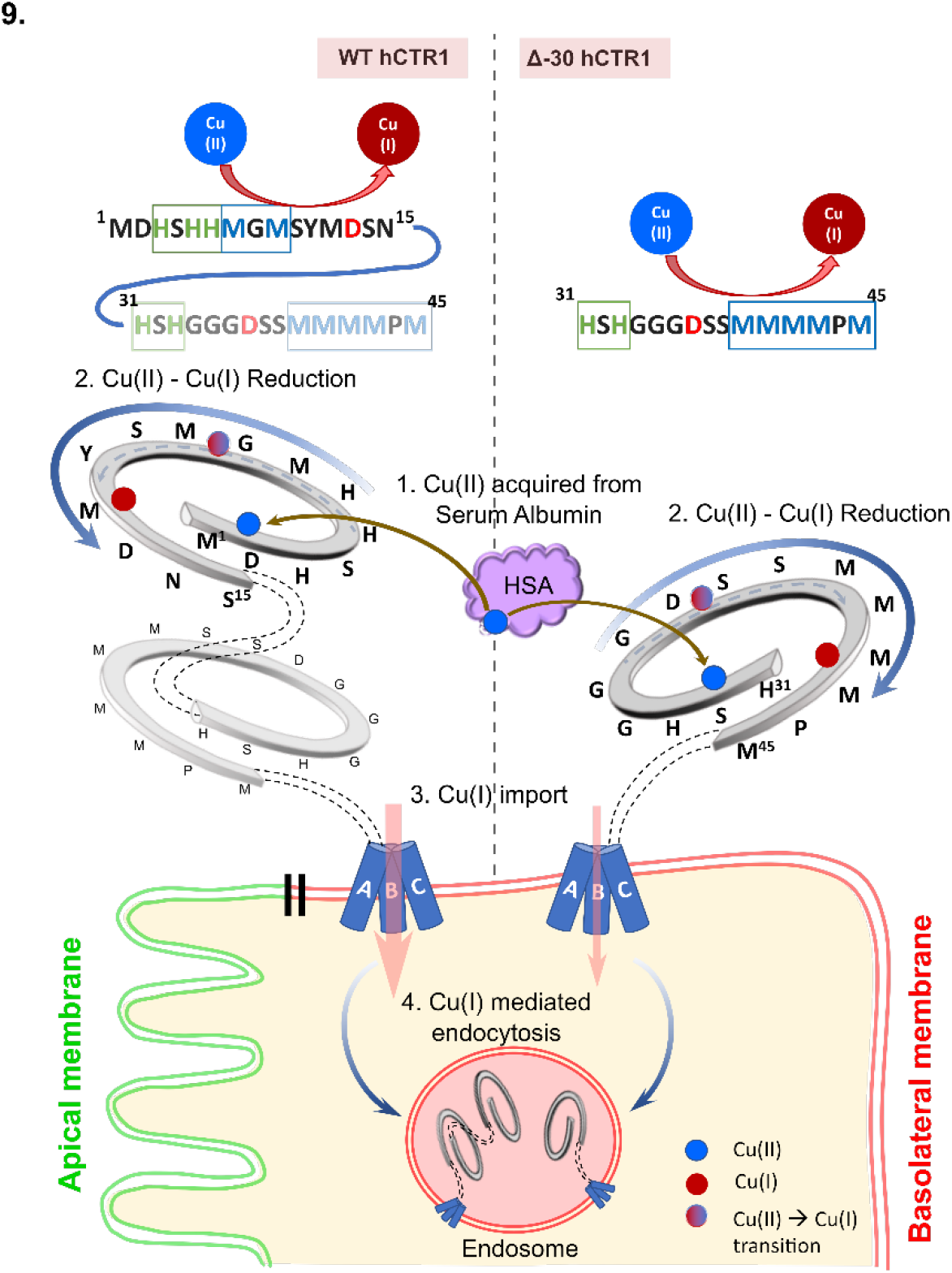
Model depicting the correlation between the major functionalities of the copper importer in both WT and Δ30-mutant constructs. The WT-hCTR1 as well as Δ30-hCTR1, residing on the basolateral membrane of polarized epithelial cells, are capable of acquiring Cu(II) from Human Serum Albumin (HSA) by their histidine-rich stretches. Subsequently, the reduction of Cu(II) to Cu(I) possibly happens in participation with reducing agents. Aspartates facilitate this reduction and mediate the transfer of copper bound to the His-rich stretches in its +2 state to the adjacent methionines, which thereby binds and stabilizes Cu(I). This leads to import of Cu(I) and finally, Cu(I) mediated endocytosis of the protein. Because of the complementary nature of His-Asp-Met clusters in the proximal and distal parts of the N-term, Δ30-hCTR1 can function like the full length WT, albeit the copper import property of the former is much lesser as compared to that of the later.

## Discussion

Copper is indispensable for the maintenance of all eukaryotic life forms. Copper exists in two redox states, Cu(I) and Cu(II). Cu(II), owing to higher stability, is more abundant in nature. Cu(I), on the other hand, though biologically relevant, has a higher tendency to get oxidized. The long conundrum that has existed in the field of biometals and, more specifically, in copper is how Cu(II) is reduced to bioavailable Cu(I). The involvement of reductases has been hypothesized in the reduction mechanism. In yeast, the FRE family of metalloreductases facilitates copper acquisition by reducing copper from cupric [Cu(II)] to cuprous [Cu(I)] state. However, no clear and tangible mechanism of this reduction has been deciphered for mammalian cells. The STEAP family proteins (STEAP-2, STEAP-3, and STEAP-4) have been shown to reduce copper *in vitro*(Ohgami et al., 2006). Over-expression of these three STEAP proteins in HEK293T cells upregulates intracellular uptake of copper. Besides STEAPs, ascorbate has also been implicated in reducing Cu(II) to Cu(I) in the blood. Another interesting phenomenon that warrants a clear understanding is how the Cu(I) stays reduced during its process of getting transported inside the cell. Using a peptide-based model, Galler *et al*. has recently shown that the trimeric arrangement of hCTR1 N-term promotes Cu(II) reduction and stabilization of the reduced form(Galler et al., 2020). However, the inference from this study has been limited by its *in-vitro* nature and the utilization of just the ATCUN motif, encompassing the first four or six residues of the N-term of hCTR1.

In this study, we have taken a combinatorial approach involving *in-silico* and *in-vivo* approaches to decipher the role of the N-term of hCTR1 in copper binding, maintaining its redox state and subsequent intracellular uptake. Further, we have delineated the relationship between copper uptake and endocytosis of the transporter.

In agreement with studies conducted by Kaplan’s and other groups, we found that hCTR1 localized on the basolateral side of polarized epithelia. We further identified the compartment as the common recycling endosome where hCTR1 endocytoses upon copper treatment. Extrapolating this finding to an organ system, the amino-terminal of the transporter would be exposed to blood proteins transporting Cu(II), e.g., albumin, under basal condition. Albumin, one of the most abundant proteins in blood serum, binds to Cu(II) and has been shown to transfer copper to the N-term of the Copper Transporter-1 in the presence of ascorbate(Du et al., 2013; Shenberger et al., 2015). We found that the N-terminal domain rich in His-Met-Asp not only binds to copper but also maintains the physiological redox state of the metal. Our model suggests that the histidine clusters in the trimeric hCTR1 bind to the Cu(II) and possibly increase its local concentration. Based upon studies from other groups, we hypothesize that ascorbate (soluble) and other membrane-bound reductases, e.g., STEAP proteins convert Cu(II) to Cu(I) on the extracellular surface. Based upon our present study, we determined that the methionine clusters in combination with aspartates on the hCTR1 N-terminal domain facilitates reduction, binds, and stabilizes the reduced copper state. Eventually, the Cu(I) enters the pore via its interaction with M150 and M154 triads situated on the entrance of the copper transporting pore. Cu(I) uptake in the cell induces endocytosis of the transporter, which regulates levels of copper import. The first methionine cluster, ^7^MGM^9^, serves as the primary coordinator for Cu(I), deletion of which leads to reduced copper uptake and subsequently abrogates hCTR1 endocytosis. Histidines, on the other hand, do not exhibit a direct interaction with Cu(I), and hence its absence has no apparent effect on copper uptake or hCTR1 endocytosis. Interestingly, in limiting but physiological copper, the role of his motifs becomes more important to increase the local concentration of Cu(II) at the amino-terminal site of the protein for its subsequent reduction. We, for the first time show the relevance of N-terminal aspartates in copper uptake and endocytosis of hCTR1. In our MD simulation model we observed that aspartates in conjunction with histidines provides key ligation for Cu(II) binding. This is the possible reason for the relative higher frequency of aspartate residues in the region coded by exon 1 as compared to the rest of the three exons.

Our study reinforces the fact that copper induced endocytosis is a self-regulatory mechanism to limit copper uptake by the transporter. We found that the mutants that show reduced copper uptake also fail to endocytose. This observation is in agreement with the previous findings that mutating the methionines of the motif ^150^MXXXM^154^ either singly or in pairs inhibits copper uptake and also exhibits defective endocytosis in response to high extracellular copper (Guo et al., 2004). We hypothesized that the copper that is reduced in the extracellular milieu and is stabilized by the methionines of the amino terminus is subsequently relayed to the ^150^MXXXM^154^ for its eventual uptake.

We observed an interesting chasm in the conservation status of the proximal and distal parts of the amino-terminal of the transporter among various eukaryotic organisms. Based upon our copper uptake and endocytic assays in human CTR1, we determined that the proximal part can complement the function of the distal part in its complete absence. Some species (like hamsters) lack the distal His-Met-Asp motifs(Soll et al., 2010). We hypothesize that the disposition of the His-Met-Asp motifs on the two regions of the amino-terminal provides flexibility among species in terms of their copper requirement and availability of copper in the environment where they thrive.

The study provides a clear understanding of the role of the amino-terminal of hCTR1 in maintaining a suitable redox state of copper that is key for its uptake. Further, we deciphered the differential participation of histidines, aspartates, and methionines in Cu(II) and Cu(I) binding. The proposed model has been illustrated in Fig 9. hCTR1 has also been implicated in the binding and uptake of the anticancer drug, cisplatin, and other platinum complexes(Song et al., 2004; Liang et al., 2014). It will be important to determine if platinum drugs imported by hCTR1 follow a similar mechanism of binding to the amino terminal, leading to its uptake and eventual endocytosis of the transporter.

## Materials and Methods

### Primers, Plasmids, and antibodies

hCTR1 was cloned in p3XFLAG CMV10 vector (Sigma #E7658, a kind gift from Dr. Rupasri Ain, CSIR-IICB) using HinDIII and EcoRI Restriction site. Amino-terminal mutations were generated by following the Q5^®^ Site-Directed Mutagenesis Kit (NEB #E0554) protocol. Primers for the site-directed mutagenesis were designed as per kit protocol and obtained from GCC biotech (West Bengal, India); their sequence details are provided below. mEGFP-ATP7B and mKO2-ATP7A constructs were available in the lab. Following are the antibodies, used for different experiments: rabbit anti-FLAG M2 (CST #14793), mouse anti-FLAG (CST #8146), anti-phalloidin 647 (Abcam #ab176759), Donkey anti-Rabbit IgG (H+L) Alexa Fluor 488 (Invitrogen #A-21206), Donkey anti-Mouse IgG (H+L) Alexa Fluor 568 (Invitrogen #A10037), Rabbit anti-GAPDH (BioBharti #BB-AB0060), goat anti-rabbit IgG-HRP conjugated secondary antibody (BioBharti #BB-SAB01A). Plasmid isolations were done by QIAGEN plasmid mini kit (QIAGEN #27104).

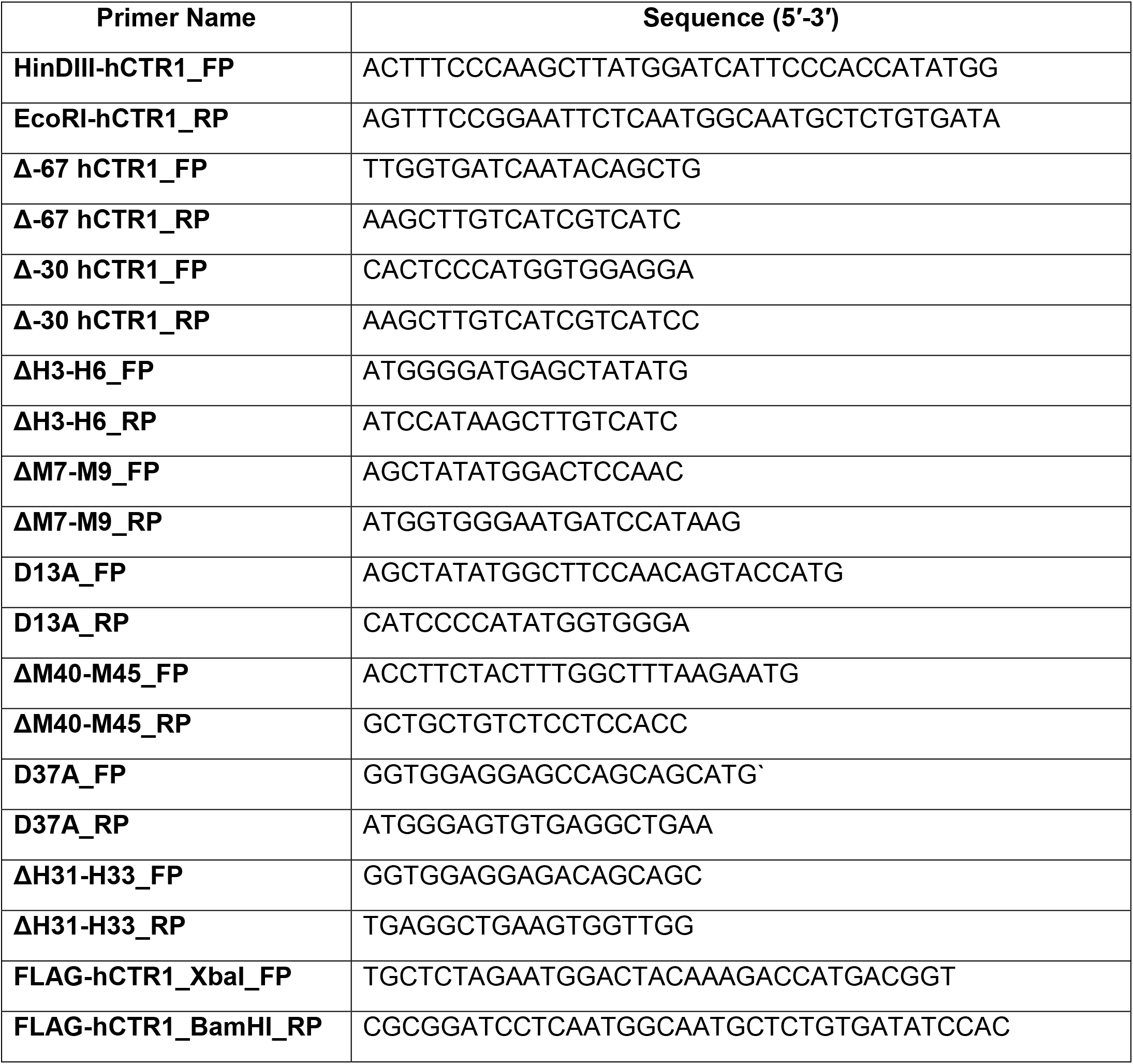

### Cell lines and cell culture

MDCK-II cells were grown and maintained in Dulbecco’s Modified Eagle’s Medium (DMEM) (Sigma #D6429) supplemented with 10% Fetal Bovine Serum (FBS, Gibco #26140079), 1X Penicillin-Streptomycin (Gibco #15140122). For transfection, electroporation was performed using Nucleofector 2b and Amaxa kit V (Programme T023). After the electroporation, 3×10^5^ cells were grown in 0.4µm inserts (Corning #3401). HEK293T cells were grown and maintained in DMEM supplemented with 10% FBS, 1X Penicillin-Streptomycin, 1X Amphotericin B (Thermo #15290026).

### Copper treatments

A stock solution (10 mM) of copper chloride (SRL #92315) dissolved in water was used as a source of Cu(II) for copper treatment. For simulating the copper-chelated condition, a stock solution (2.5 mM) of TTM (Sigma #323446) dissolved in DMSO (Sigma #D2650) has been used. Direct Cu(I) was provided to the cells by treating with a mixture of CuCl2, L-ascorbic acid (Sigma-Aldrich #95209) and THPTA in ratio 1:2:2 in HBSS (Gibco #14025092). The later two components were also dissolved in water and a stock solution of 10 mM concentration each was prepared. All treatments were applied only on the baso-lateral side of the cells, if not mentioned otherwise.

### Transferrin internalization assay

For 633-Tf uptake assays, cells were starved for 60 min at 37^0^C in HBSS containing 20 mM HEPES (standard buffer) and incubated for 60 min at 4^0^C with 10 mg/ml 633-Tf in 1% BSA standard buffer. After that, temperature was shifted to 37^0^C and, after the specified time, cells were immediately rinsed (with ice-cold HBSS) and fixed (ice-cold 4% PFA in PBS). After quenching PFA (50 mM NH4Cl in PBS) for 15 min, cells were permeabilized with 0.01% triton X-100 in PBS for 7 min (Perez Bay et al., 2013). Following this the cells are incubated with the corresponding primary antibodies for 2 hrs and then secondary antibodies again for 2 hrs in 1% BSA in PBS, and washed three times after each incubation.

To label BSE(basal sorting endosome) and CRE(common recycling endosome), Tf-633 was added to the basolateral side of the cells, growing on the membrane, for 5 min and 30 min respectively at 37^0^C (25 ug/ml in 1% BSA standard buffer) (Perez Bay et al., 2016).

### Immunofluorescence and microscopy

All the above-mentioned treatments were done after the cells reached polarization. After washing with ice-cold PBS (2×2 mins), cells were Fixed with 2% PFA in PBS for 20 mins at room temperature (RT), followed by 20 mins incubation with 50 mM ammonium chloride in PBS for quenching extra PFA. Next the cells were washed with PBS and blocking was performed in 1% Bovine Serum Albumin (BSA, SRL #85171) in PBSS (0.075% saponin in PBS) for 20 mins at RT. Primary antibody incubation was performed for 2 hrs at RT followed by PBSS washes (3×5 mins). After that, incubation with the respective secondary antibodies were done for 2 hrs followed by 5 PBSS washes. The membrane was mounted with the Fluoroshield with DAPI mountant (Sigma #F6057). All images were acquired with Leica SP8 confocal platform using oil immersion 63X objective (NA 1.4) and deconvoluted using Leica Lightning software.

### Image analysis

Images were analyzed in batches using ImageJ (Schneider et al., 2012), image analysis software. For colocalization study, the Colocalization_Finder plugin was used. ROIs were drawn manually on the best z-stack for each cell. Manders’ colocalization coefficient (MCC) (MANDERS et al., 1993) was used for quantifying colocalization. Macro used in ImageJ are available in https://github.com/saps018/hCTR1-N-term/tree/main/colocalization.

### Statistical analysis

For statistical analysis and plotting, ggplot2 (Wickham, 2009) and ggpubr (Kassambara, 2019) package was used in R v-4.0.4 (Team, 2013). Non-parametric tests for unpaired datasets (Mann-Whitney U test) were performed for all the samples.

### Determination of cellular copper concentrations by ICP-OES

After the differential treatments have been applied to the polarized MDCK-II cells, they were washed with HBSS in a cold chamber and then incubated with TryplE Express (Gibco #12605028). Cells were harvested by scraping. Cells were pelleted down at 2,500 rpm for 3 mins and were washed 5 times with ice-cold DPBS (Gibco #14200075). The pellets were finally dissolved in DPBS and were counted by haemocytometer using Trypan Blue stain (Gibco #15250061). Further experiments were carried out with 2.5×10^6^ cells for each condition. Cell samples were digested for 16 hours with 100 µL 65 % ICP-OES grade HNO_3_ at 95^0^C. After digestion, samples were diluted in 5 mL of double distilled water and were syringe filtered through a 0.45-micron filter. Copper calibration is done by acid digestion of copper foil (procured from Alfa Aeasar) in 10 mL suprapure HNO_3_ for 1 h. (MWD conditions: Power=400 W; Temperature=100^0^C; Hold time= 1 h). From the obtained solution, different solutions of varying copper strengths (50, 75, 100, 250, 500, 1000, 5000, 10000 ppb) were prepared and were used for calibration. Copper concentration was determined using a Thermo Scientific Inductively coupled plasma optical emission spectroscopy (ICP-OES) iCAP 6500.

### Strains, media, growth conditions for yeast complementation assay

For yeast complementation studies, *Saccharomyces cerevisiae* BY4742 (wild-type) strain (MATα his3Δ1 leu2Δ0 lys2Δ0 ura3Δ0) and an *S. cerevisiae* strain carrying an yCTR1 deletion (ΔyCTR1) in the BY4742 background purchased from Euroscarf (Oberursel, Germany) were used. YPD (yeast extract, peptone, and dextrose) medium was used for routinely maintaining both wild-type and deletion strains. For complementation assay, synthetic defined (SD) minimal media containing YNB (yeast nitrogen base), ammonium sulfate, and dextrose supplemented with histidine, leucine, lysine, and methionine (80 mg/liter each) was used. Yeast transformations were carried out using the lithium acetate method (Gietz and Woods, 2002). Human CTR1 was cloned in the yeast expression vector, p416TEF (as a positive control) and amino-terminal mutants were generated in the hCTR1-p416TEF construct by site-directed mutagenesis using Q5^®^ Site-Directed Mutagenesis Kit (NEB #E0554) protocol. The p416TEF vector contains a URA3 selection marker allowing growth in absence of uracil. Wild-type strain was transformed with an empty vector to allow its growth on SD-Ura (SD medium without uracil). Yeast transformants were selected and maintained on SD-Ura at 30°C.

### In Vivo Functional Complementation assay in *S. cerevisiae* by dilution spotting

Yeast transformants were grown overnight at 30°C with shaking at 200 rpm in SD-Ura medium. The primary culture was used to inoculate secondary culture in the same selective medium and was allowed to grow at 30°C till OD_600_ reached about 0.6. The cells were centrifuged, washed, and diluted in sterile water at OD_600_ = 0.2. Serial dilutions were then made with sterile water (OD_600_ = 0.2, 0.02, 0.002, 0.0002), and 10μl of cell suspension from each were spotted on plates containing yeast extract, peptone, ethanol, and glycerol (YPEG plates). Plates were incubated at 30°C for 3-5 days and photographs were taken by Chemi Doc (BioRad).

### MD simulation

#### a. System Setup for Simulation

The starting structure of the hCTR1 protein is an electron crystallography structure provided by Professor Vinzenz Unger, Department of Molecular Biosciences, NorthWestern University (Aller and Unger, 2006). Only the extracellular N-terminal domain of the three monomers of the protein (residues 1 to 67) are considered for the simulations (system setup shown in Fig S3A). The protein and water are represented using the CHARMM36 force field (MacKerell et al., 2001). The Cu(II)/Cu(I) are represented as virtual site models in an octahedral and tetrahedral geometry, respectively (Liao et al., 2015; Pang, 1999). Each of the Nterm-copper systems was solvated by ∼16000 TIP4P water molecules in a box of dimensions 80 x 80 x 80 Å3. The physiological concentration (150 mM) of Na+ and Cl-ions along with extra Na+ ions were used to neutralize the system. Simulations were performed using molecular dynamics software GROMACS 2019.6 (GROMACS development team, 2019).

#### b. Equilibration and Simulation

Initially each system is energy minimized using the steepest descent method (Fritsch et al., 1988) for 10000 steps, followed by heating it to 300K in 200 ps using Berendsen thermostat and barostat(Berendsen et al., 1984) with a coupling constant of 0.5 ps each. Restraints of 25 kcal/mol/Å^2^ are applied on heavy atoms during the heating process. Thereafter, equilibration is carried out for 2 ns at constant temperature (300 K) and pressure (1 bar) without any restraints using the same thermostat and barostat with coupling constants of 0.2 ps each. The last 100 ps of NPT simulation is used to calculate the average volume which is used in all simulations going forward. Unrestrained NVT equilibration for 200ns at temperature 300K is carried out using the velocity-rescale thermostat (Bussi et al., 2009) with coupling constant of 0.1 ps. During the simulation, LINCS algorithm (Hess et al., 1997) is used to constrain all the bonds and Particle Mesh Ewald (PME) method (Darden et al., 1993) is used for electrostatics. The distance cut-offs for the van der Waals (vdW) and electrostatic long-range interaction is kept at 12 Å. The time step for each simulation is taken to be 1 fs.

#### c. Free energy calculation using Metadynamics

The equilibrated N-terminal domain is initially simulated for 5 ns in the presence of an unbound copper ion. Free-energy calculations are performed after the Cu(I)/Cu(II) binds to the N-terminal domain. To calculate the binding free energy of the ion in both of its oxidation states, well-tempered metadynamics (Barducci et al., 2008) simulations are performed after equilibration using distvec (Chowdhury et al., 2021) (Fig S3B) and native contacts (N_c_) (Fig S3C) as collective variables. We performed a long (∼150 ns). metadynamics simulation with a hill height of 0.2 kJ/mol and a bias factor of 10, and hills deposition rate of 2 ps. Gaussian widths for distvec and native contacts are taken to be 0.6 Å and 5, respectively. An upper wall restraint is applied at 45 degrees on the angle between two vectors and, as shown in Figure S3B of SI. For free-energy calculations, PLUMED 2.6. (Tribello et al., 2014) is used along with GROMACS. The system size and run lengths of all the systems are provided in the Table S3D of SI.

### Synthesis of Tris-(3-hydroxypropyl triazolyl methyl)-amine (THPTA)

The synthesis and characterization of THPTA has been described in detail in this paper(Hong et al., 2009).

### Electron Paramagnetic Resonance

Electron paramagnetic resonance (EPR) data were recorded with an EMX MICRO X, Bruker spectrometer, operating at a microwave frequency of approximately 9.75 GHz. Spectra were recorded using a microwave radiation power of 10 mW across a sweep width of 2000 G (centred at 2200 G) with modulation amplitude of 10 G. Experiments were carried out at 100 K using a liquid nitrogen cryostat.

EPR samples were prepared from 5mM CuCl_2_, 10mM ascorbate and 10 mM THPTA solutions dissolved in water in order to distinguish between the oxidation states of copper in presence of ascorbate and/or THPTA. Samples were frozen in a quartz tube after addition of 10 % glycerol as a cryoprotectant and stored in liquid nitrogen until used.

### UV-Visible spectroscopy

UV-Vis spectra were recorded at room temperature on Cary 8454 Agilent spectrophotometer, over the spectral range 300–800 nm, using the 1 cm-path-length quartz cuvettes. 20mM CuCl_2_ were mixed with different equivalents (ranging from 0.5 to 2 equivalence) of ascorbate and 2 equivalences of THPTA in water and spectra were recorded at two different time points, 0 min and 30 mins.

### Sequence alignment and conservation status

CTR1 sequences all over the species were obtained from NCBI. Exon composition of chordate CTR1 was obtained from Ensembl (https://www.ensembl.org/)(Yates et al., 2020). Sequence alignment file was created in Clustal Omega(Sievers and Higgins, 2014; Sievers et al., 2011). For sequence visualization, Jalview 2.11.1.4 was used(Waterhouse et al., 2009). Amino acid composition details were created using python script. Protter was used for preparing a representative image of CTR1(Omasits et al., 2014). Python codes used in exon analysis are available in https://github.com/saps018/hCTR1-N-term/tree/main/Exon%20analysis.

## Acknowledgments

This work is supported by DBT-Wellcome Trust India Alliance Fellowship (IA/I/16/1/502369), Early Career Research Award (ECR/2015/000220) from SERB, Department of Science and Technology (DST), Government of India, and IISER-K intramural funding to AG. SK, RP, SM, BM are supported by a Pre-doctoral fellowship from Council of Scientific and Industrial Research (CSIR), India. The Pre-doctoral fellowship for Ruturaj is supported by Intramural Institute funding (IISER-K). SS is supported by KVPY fellowships from the Government of India. We thank Prof. Anand K Bachhawat (IISER-Mohali) for helping us with the technicalities of the yeast experiment and Ms. Siddhanta Nikte, Ph.D. student from CSIR-NCL, for help and discussions regarding the simulations and metadynamics calculations. We would also like to thank the ICP-OES facility, IISER-Kolkata, for helping us with our experiments. We would like to acknowledge Dr. Ashima Bhattacharjee (Amity Univ. Kolkata), Prof. Arnab Mukherjee (IISER Pune), Rituparno Chowdhury (IISER Kolkata) and Dr. Sayam Sengupta (DCS, IISER-Kolkata) for providing critical review of the manuscript.

## Author contributions

Conceptualization: AG,SK,SS,SM; Methodology: AG,SK,SS,SM,RR,RP,SD; Software: SM,SS,SK; Validation: AG,SK,SM,SS,DS; Data analysis: AG,SM,SK,SS; Investigation: AG,SK,SM,RP,DS; Resources: AG,SK,SS,RR,RP,BM,ERB,RS; Writing and editing: all authors.

**The authors declare no competing financial interests.**

---

**S.1.**
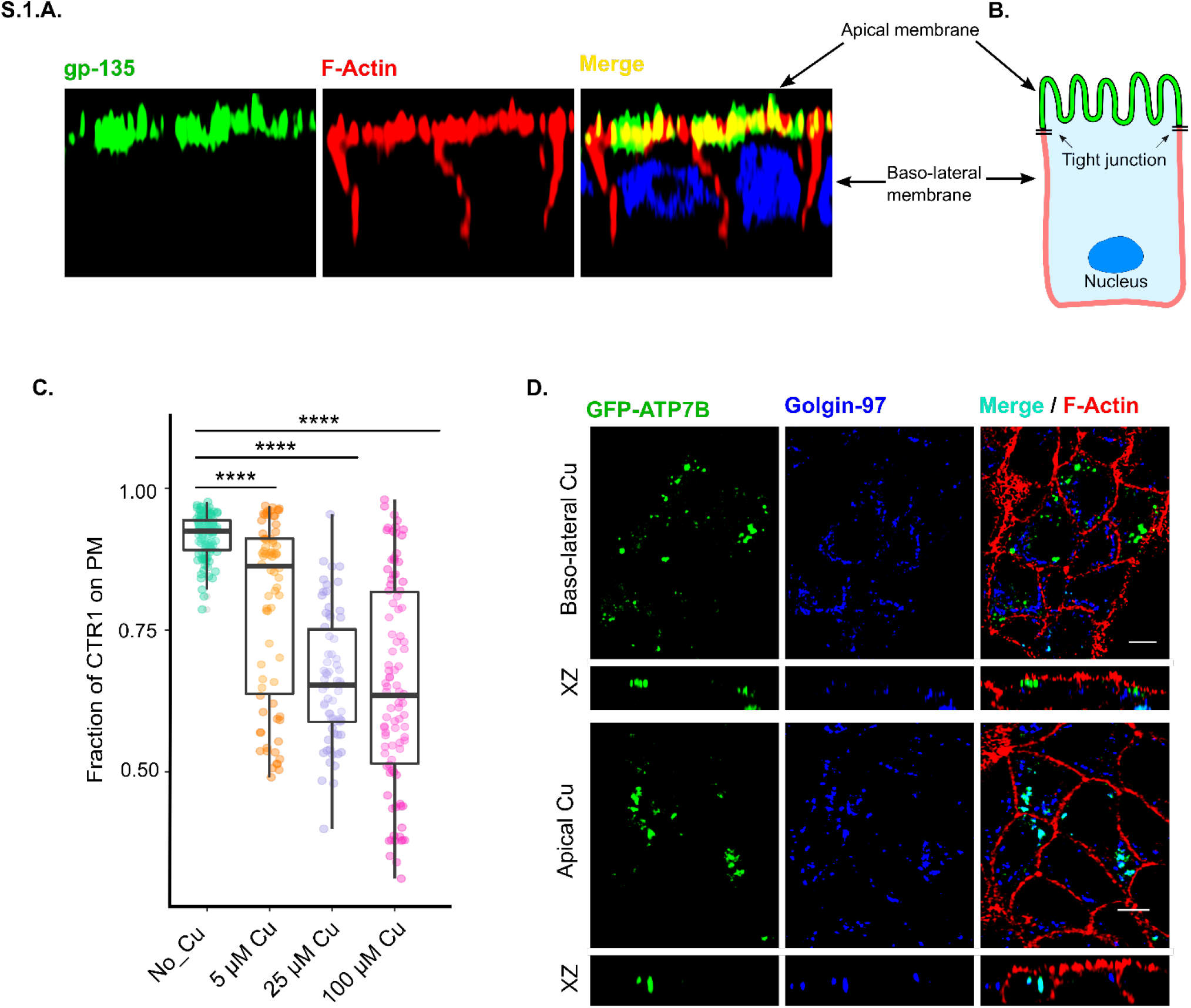
**(A)** Representative image of polarized MDCK-II cell line after fixation and staining. (Upperside is the Apical side, stained with gp-135 (also known as podocalyxin); (Green-gp-135, Red-F-Actin, Blue-DAPI). After the polarization, cells attain height ∼ 12-15 µm and form distinct apical and basolateral sides separated by gap junctions. (**B).** Representation of a polarized MDCK-II cell. (**C).** Fraction of WT-hCTR1 colocalization with membrane marker F-actin, demonstrated by box plot with jitter points. The box represents the 25–75th percentiles, and the median in the middle. The whiskers show the data points within the range of 1.5× interquartile range (IQR) from the 1st and 3rd quartile. ****P<0.0001 (non-parametric Mann–Whitney U test/Wilcoxon rank-sum test). Sample size (n) for Basal: 100, 5 uM Cu: 74, 25 uM Cu: 66, 100 uM Cu: 101. (**D).**Transiently transfected GFP-ATP7B shows trafficking towards the apical membrane (upper panel) upon copper treatment in the basolateral chamber whereas the protein shows no trafficking from Golgi during copper treatment only on the apical side (lower panel) [In all the conditions, cells are polarized MDCK-II, XZ section shows the orthogonal sections of all the stacks, green-GFP-ATP7B, blue-Golgin-97 and red-F-actin; Cu treatment-100μM; scale bar-5μm].

**S.2.**
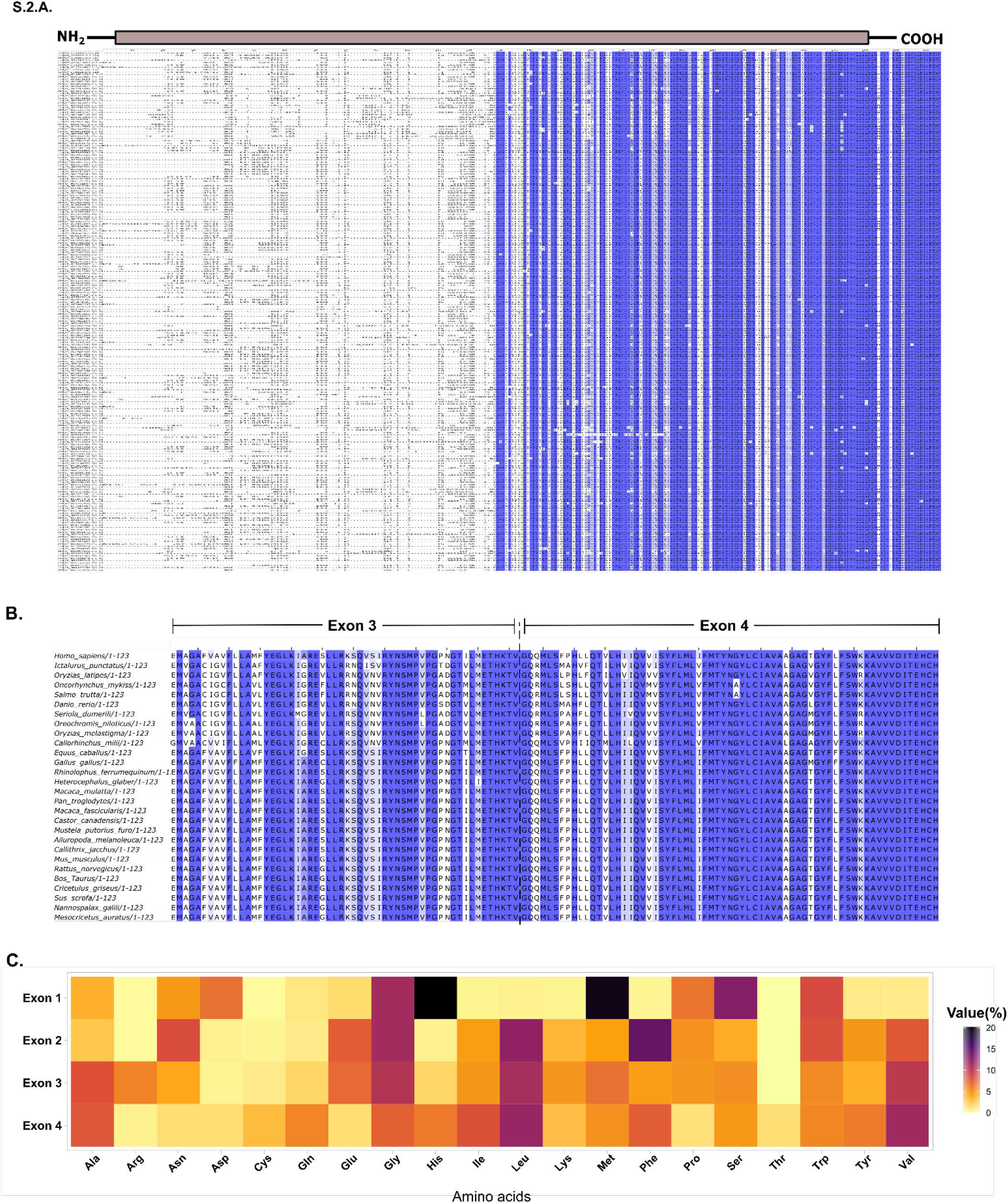
Sequence alignment of CTR1s across the species **(A)** and last two exons of chordata CTR1s **(B)**. Conserved residues are marked by blue colour. The intensity of the blue colour is proportional to the conservation status of a residue. **(C)** Heat map showing the percentages of the different amino acids encoded by each of the four chordate CTR1 exons.

**S.3.**
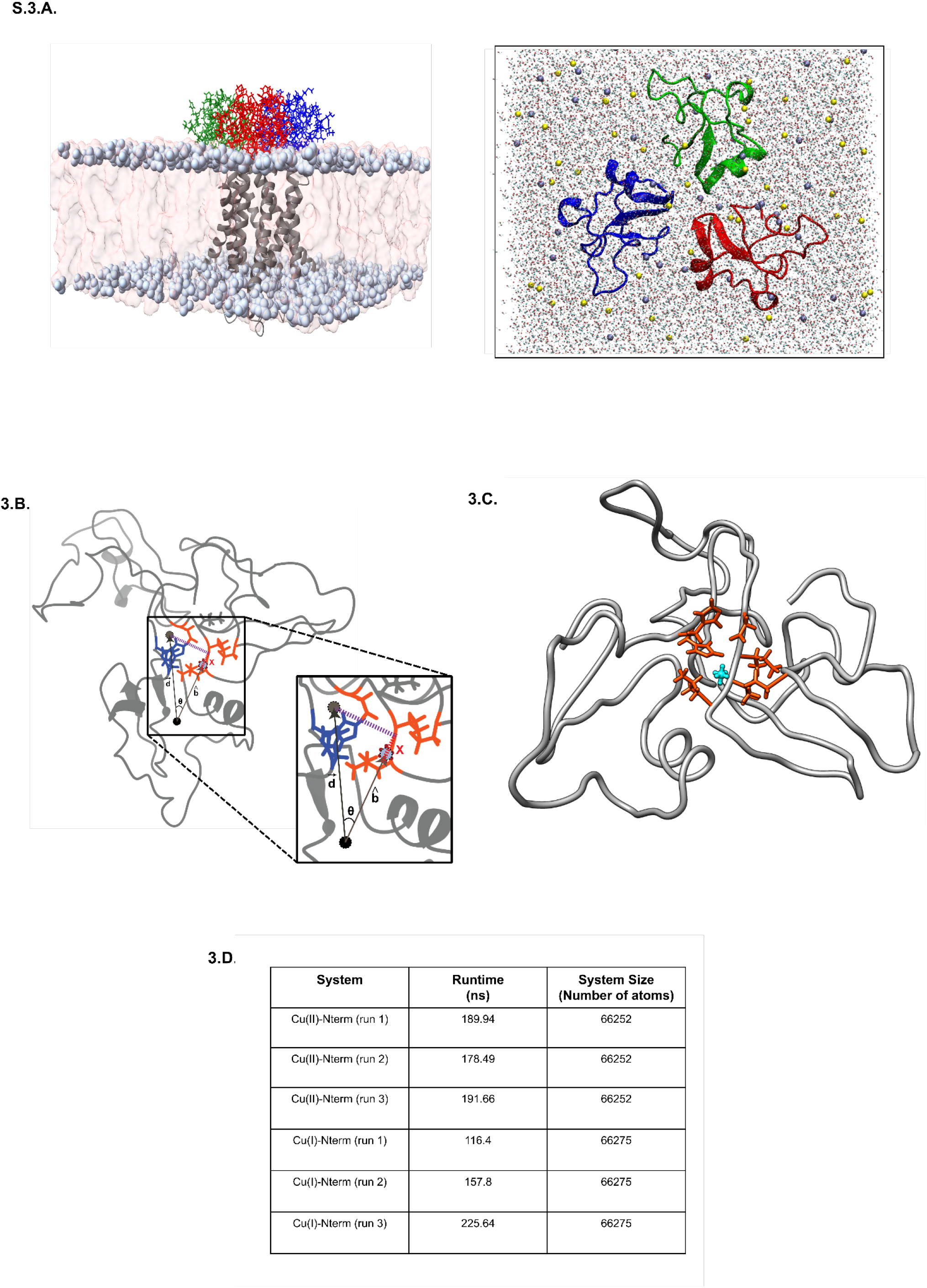
**(A)** hCTR1 is a trimeric membrane protein (transmembrane regions depicted as ribbons in grey) that is shown to be embedded in a POPC lipid bilayer (shown as a surface with phosphate head groups depicted as spheres) (left panel). The simulation setup in this work comprises only the extracellular N-terminal region of the protein in water. Na^+^ ions are shown in yellow and Cl^-^ ions in mauve. [top view of the system in (right panel)]. The trimeric conformation of the N-terms is maintained by putting position restraints on the heavy atoms of the last residues of each monomer. **(B)** The schematic description of the distvec collective variable which is mathematically specified as the body-fixed distance to the ligand/ion of interest. In the figure the protein is shown as a grey silhouette, the c.o.m of the protein is shown by the black circle, the c.o.m of the binding region is given by the violet circle, and the c.o.m of the Cu(II)-octahedral dummy model is depicted by the light blue circle. The residues marked in deep blue and orange are the histidine-rich and histidine-deficient halves of the binding site respectively, which are used to body-fix the coordinate axes **(C)** The Native contact (***NNcc***) is defined by the spatial proximity of groups of atoms ***ggAA*** and ***ggBB*** in the native state. The Cu(II) virtual site model (shown in cyan) constitutes ***ggAA*** while the heavy atoms of part of the protein that resides within 5.5 Å from the c.o.m of the copper dummy model constitute ***ggBB*** (shown in orange). **(D)** The system size and simulation lengths for the metadynamics simulation for all the systems are given in **Table 3D**.

**S.4.**
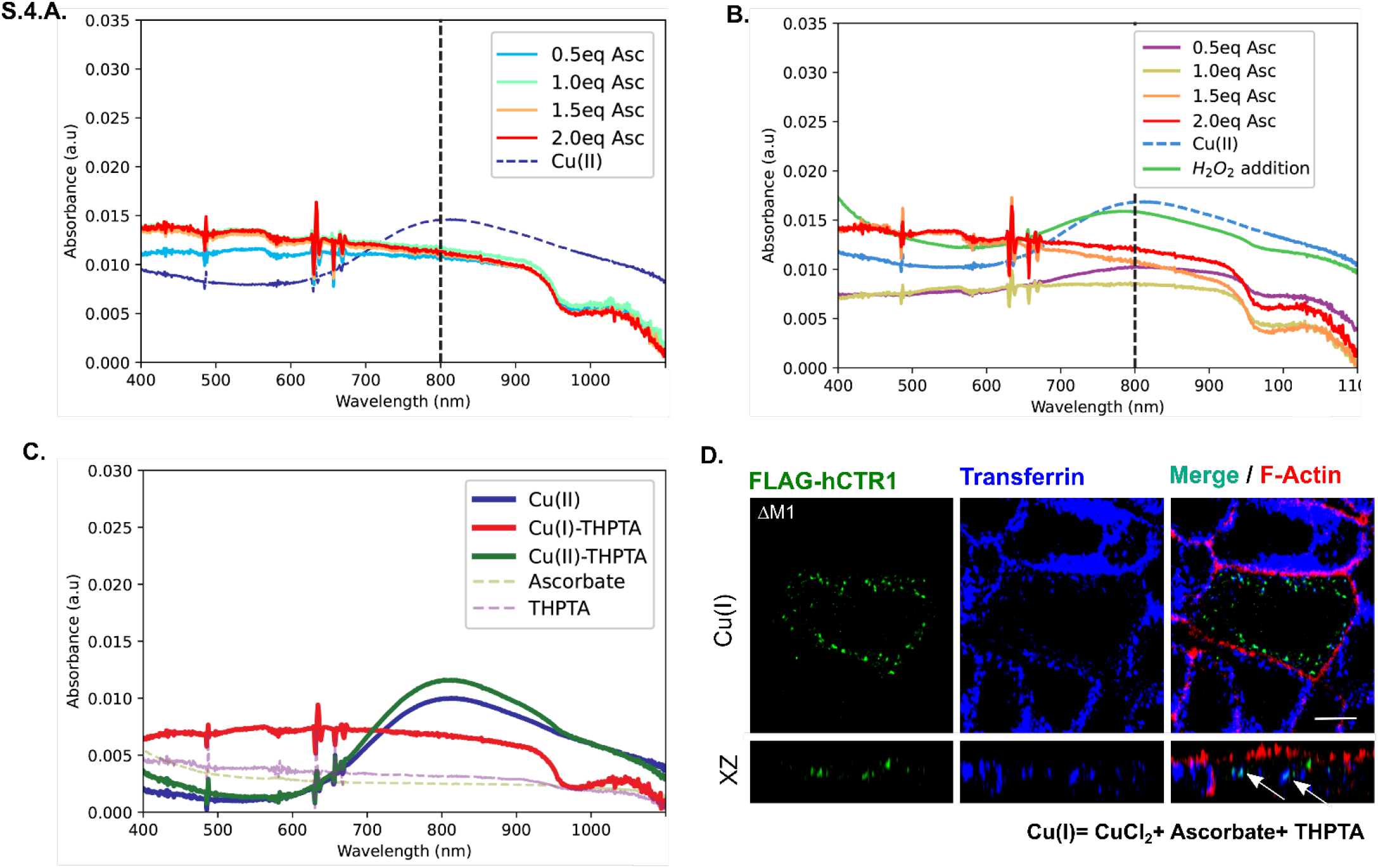
UV-Vis Spectra shows a wide band for Cu(II) with an absorption maximum at 800 nm. 20mM CuCl_2_ in water is treated with 2 equivalence of THPTA and varying equivalence of ascorbate ranging from 0.5 to 2 equivalence at two different time points, 0 min **(A**) and 30 min **(B)** respectively. The peak for Cu(II) is absent in all 4 situations at both the time points, however, it was decided to use 2 equivalence of ascorbate for our subsequent *in-vivo* experiments, just to ensure that copper remains in its reduced state during the entire duration of our experiments. **(B)** Addition of an oxidising agent, H_2_O_2_ to the ascorbate+THPTA+CuCl_2_ (2:2:1) mixture, at the end of 30 mins causes the peak for Cu(II) to return, indicating the reoxidation of Cu(I). **(C)** The peak for Cu(II) at 800 nm retains itself when 40mM THPTA is added to 20mM CuCl_2_ solution (Cu(II)-THPTA), but in presence of both 40mM ascorbate and 40mM THPTA, the peak vanishes indicating the formation of Cu(I)-THPTA, which is not UV-Vis active (**D).** ΔM1 (ΔM7-M9) endocytoses when treated with ascorbate +THPTA + CuCl_2_ in 2:2:1 ratio {readymade source of Cu(I)} and colocalizes with the basolateral sorting endosomes and common recycling endosomes (marked by post-30 min internalization of Transferrin-Alexa 633). [In this condition, the cell is polarized MDCK-II, XZ section shows the orthogonal sections of all the stacks, green-FLAG-hCTR1, Blue-Transferrin and red-F-actin; (200μM ascorbate + 200μM THPTA + 100μM CuCl_2_) treatment on the basolateral chamber of the transwell, scale bar-5 μm].

**S.5.**
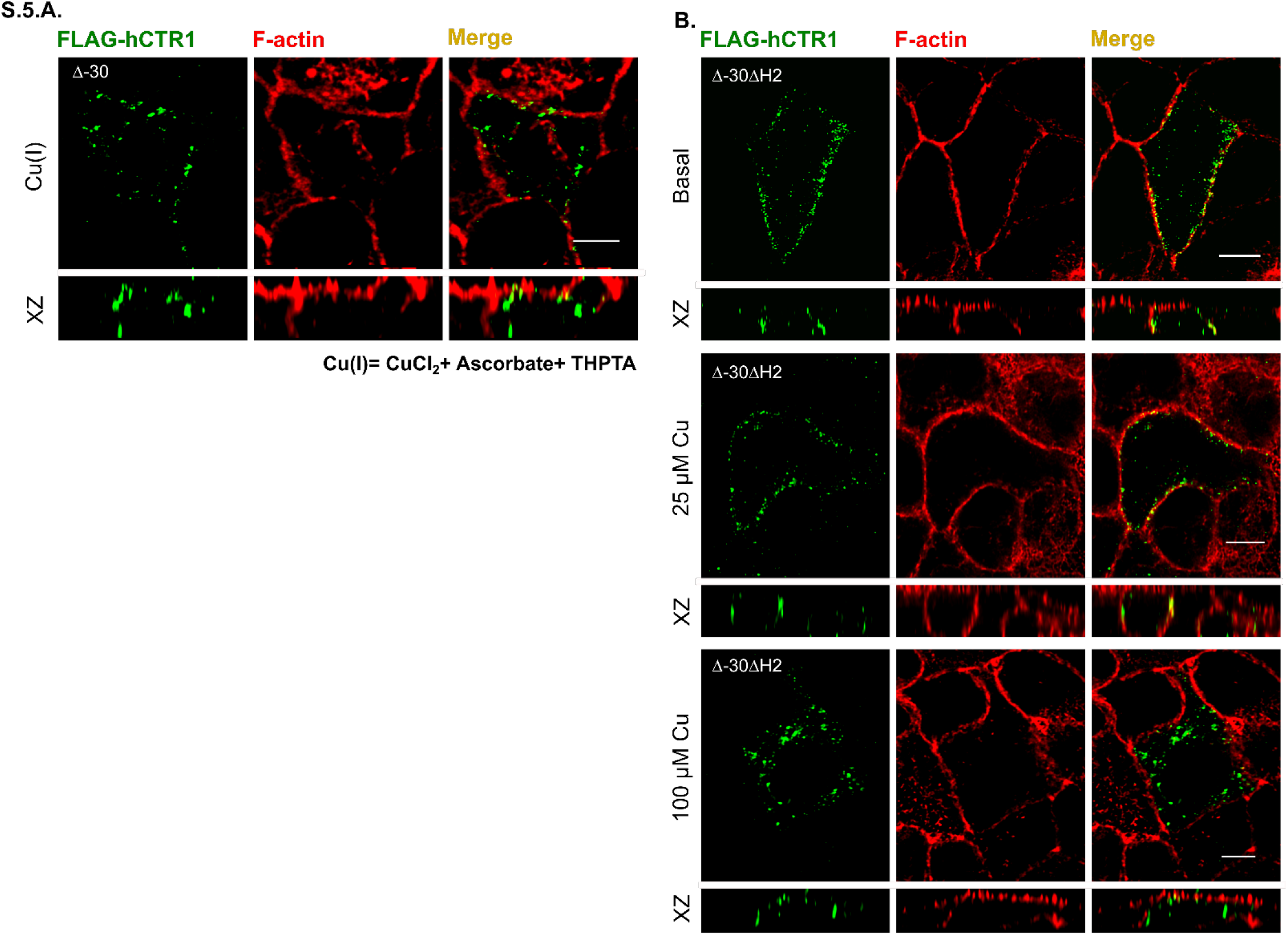
**(A)** Δ-30 exhibits endocytosis when treated with the ready source of Cu(I). (**B).** Δ-30ΔH2 mutant resides on the PM under basal (upper panel) conditions, 25μM Cu treatment (middle row) also fails to induce endocytosis of the same. In response to 100μM Cu (lower panel) however, Δ-30ΔH2 endocytoses. [In all the conditions, cells are polarized MDCK-II, XZ section shows the orthogonal sections of all the stacks, green-FLAG-hCTR1 and red-F-actin; treatments are on the basolateral chamber of the transwell, scale bar-5 μm].

